# Pluripotent Stem Cell Plasticity is Sculpted by a Slit-Independent Robo Pathway in a Regenerative Animal

**DOI:** 10.1101/2025.04.14.648795

**Authors:** Kuang-Tse Wang, Yu-Chia Chen, Fu-Yu Tsai, Catherine P Judy, Carolyn E Adler

**Affiliations:** Department of Molecular Medicine, Cornell University College of Veterinary Medicine, Ithaca, NY, USA; Department of Genetics, Harvard Medical School, Boston, MA, USA; Department of Orthopedic Research, Boston Children’s Hospital, Boston, MA, USA

**Keywords:** Stem cell plasticity, *roboA*, Anosmin-1, *foxA*, planarian regeneration, neural differentiation, Kallmann syndrome

## Abstract

Whole-body regeneration requires adult stem cells with high plasticity to differentiate into missing cell types. Planarians possess a unique configuration of organs embedded in a vast pool of pluripotent stem cells. How stem cells integrate positional information with discrete fates remains unknown. Here, we use the planarian pharynx to define the cell fates that depend on the pioneer transcription factor FoxA. We find that Roundabout receptor RoboA suppresses aberrant pharynx cell fates by altering *foxA* expression, independent of the canonical ligand Slit. An RNAi screen for extracellular proteins identifies Anosmin-1 as a potential partner of RoboA. Perturbing global patterning demonstrates that *roboA*/*anosmin-1* functions locally in the brain. By contrast, altering pharynx fate with *foxA* knockdown induces head-specific neurons in the pharynx, indicating a latent plasticity of stem cells. Our data links critical extracellular cues with cell fate decisions of highly plastic stem cells, ensuring the fidelity of organ regeneration.

## Introduction

Regeneration is defined by the ability of tissues to replenish specific cell types in discrete anatomical positions, ensuring both structural integrity and functional restoration. This process often relies on stem cells to sense surrounding tissues, which guides subsequent replacement of necessary cell types. Because injuries are inherently unpredictable, stem cell plasticity is thought to contribute to successful regeneration by integrating extrinsic cues with downstream transcriptional outputs ^1–3^. Despite its importance, the molecular mechanisms governing these interactions during regeneration remain elusive.

The planarian flatworm *Schmidtea mediterranea* is a classical model organism for studying whole-body regeneration ^4^. This remarkable ability is driven by a population of pluripotent stem cells capable of giving rise to all cell types throughout the animal’s life ^5,6^. These stem cells are distributed across the body, while individual organs are spatially restricted. Whether the differentiation potential of stem cells varies with body position and how extracellular signals ensure organogenesis at the proper location are unknown ^3^.

Two systems that broadly influence the maintenance of organ positioning and identity have been previously proposed. ‘Position control genes’ (PCGs), such as *wnt* and *bmp*, are regionally expressed by muscle cells ^7^. Their knockdown causes strong disruptions in axis patterning and duplications of organs ^8–10^, similar to homeotic transformations during development ^11^. Although these findings highlight potential extracellular cues that define broad tissue boundaries, how stem cells receive these signals and adopt specific fates in the vicinity of organs is an open question. The ‘target zone’ model proposes that pre-existing organs can direct stem cell differentiation toward appropriate cell types ^1,12,13^, thus maintaining organ positioning and structure. Although both systems suggest that extracellular cues regulate cell differentiation, the identity of these signals and how stem cells interpret them to adopt specific fates remain largely unclear.

The pharynx is an organ with a unique anatomical position and known stem cell origin ^14,15^. It resides in a pouch in the middle of the animal body and consists of epithelium, muscles, and a radially-symmetric nerve net ^14–16^. Previous studies have demonstrated that the Forkhead transcription factor FoxA is essential for pharynx regeneration and the recovery of feeding behavior ^17,18^. Pharynx regeneration initiates from a subset of stem cells that express *foxA* transcripts, scattered around the pharynx ^19^. Together, these characteristics establish the pharynx as an excellent model for studying stem cell plasticity, where the stem cells differentiate into multiple cell types to restore a complex organ at a stereotypical position.

Prior work has identified genes responsible for inducing an abnormal pharynx either in the wrong position ^8,10^ or with disrupted anatomy ^20^. In particular, knockdown of the Roundabout receptor RoboA induces ectopic but nonfunctional pharynges ^21^. Although Robo is best known for its roles in patterning the nervous system during development, it also regulates stem cell differentiation and tissue patterning in diverse model organisms ^22–25^.

In this study, we capitalize on the *roboA*(RNAi) phenotype to explore the mechanisms that guide spatial restriction of stem cell fates to maintain organ identity. We characterize pharynx-specific lineages and find that *roboA*(RNAi) induces ectopic pharynx neurons (EPNs) and muscles in the brain of homeostatic animals, independently of *slit* and neural patterning. Further, we identify a homolog of the secreted protein Anosmin-1 (Anos1) as a potential signaling partner of RoboA. The pioneer factor FoxA is the primary determinant that toggles these fates within stem cells. This RoboA/Anos1/FoxA triad acts downstream of global patterning. Taken together, our findings demonstrate that stem cells exhibit a latent plasticity that is suppressed by RoboA and Anos1 to reinforce organ identity mediated by the downstream transcription factor FoxA.

## Results

### *roboA*(RNAi) induces ectopic pharynx cells in the head

The single planarian pharynx is maintained by stem cells expressing the transcription factor *foxA* that are distributed in the surrounding parenchyma (Figure 1A) ^26^. How *foxA*^+^ stem cells and the pharynx are restricted to this region is unclear. Previous work showed that knockdown of *roboA* induces animals to regenerate supernumerary pharynges upon amputation ^21^. We hypothesized that this atypical anatomy might provide an inroad to elucidate the underlying mechanisms that normally confine pharynx fate and position. To understand which cell types in the pharynx arise abnormally in *roboA*(RNAi) animals, we analyzed the distribution of pharynx cells (see Supplementary Text and Figure S1) by *in situ* hybridization (ISH).

**Figure 1.**
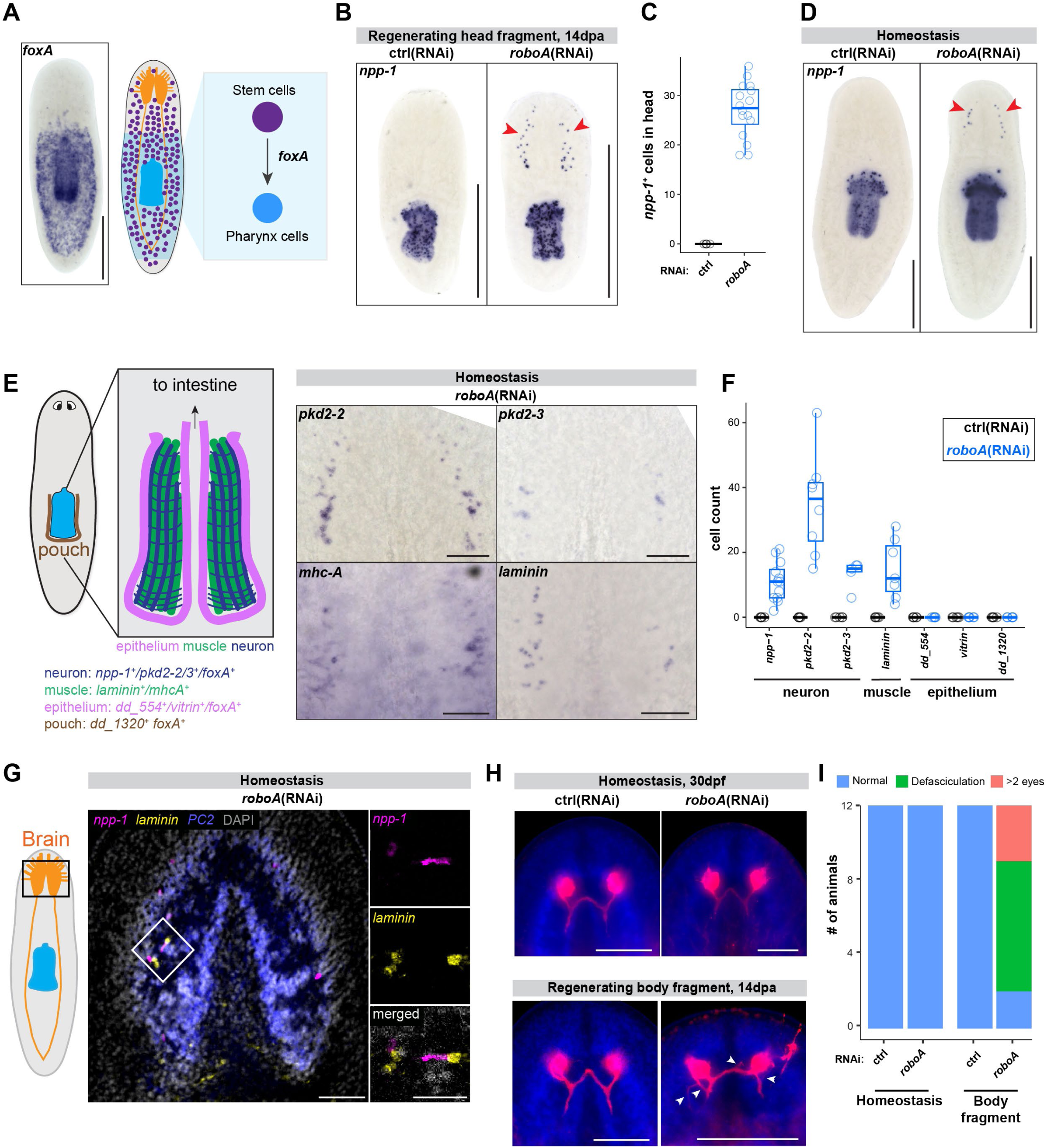
*roboA* suppresses formation of EPNs and pharynx muscles in the brain. (A) Whole-mount *in situ* hybridization (ISH) of *foxA*. *foxA* is expressed in stem cells and pharynx cells; schematic of stem cell lineage is on the right. (B) ISH of pharynx neuron marker *npp-1* in regenerating head fragments 14 days post-amputation (dpa). *npp-1* expression is restricted to the pharynx in control animals; ectopic pharynx neurons (EPNs) emerge in the head of *roboA*(RNAi) animals (red arrowheads). Scale bar = 500 μm. (C) Quantification of EPNs in head fragments at 14 dpa. (D) ISH of *npp-1* showing EPNs (red arrowheads) in homeostatic RNAi animals, 14 days post-last RNAi feed-ing (dpf). Scale bar = 250 μm. (E) Left, schematic of the cell types in the pharynx with unique markers listed below. Right, ISH of phar-ynx-specific neuron markers (*pkd2-2* and *pkd2-3*) and muscle markers (*mhc-A* and *laminin*) in the brain regions of homeostatic *roboA*(RNAi) animals, 14dpf. Scale bar = 100 μm. (F) Quantification of ectopic pharynx-specific cells in the brain regions of homeostatic *roboA*(RNAi) and control animals. (G) Confocal images of the brain of a homeostatic *roboA*(RNAi) animal stained with hybridization chain reaction (HCR) for *npp-1* (magenta), *laminin* (yellow) and *prohormone convertase* (*pc2,* blue). Insets are zooms of the white box. Scale bar = 100 µm; zoomed region = 25µm. (H) Immunostaining for Arrestin (red) in homeostatic animals (30 dpf) and body fragments with regenerated heads (14 dpa). White arrowheads highlight defasciculation. Scale bar = 200 μm. (I) Quantification of phenotypes from (H).

To determine whether pharynx number correlated with the severity of the injury, we varied the anterior-posterior amputation plane (Figure S2A). Using *neuropeptide precursor-1* (*npp-1*) as a highly specific marker of pharynx neurons ^27^, we found that the number of ectopic pharynges increased when the amputation occurred closer to the pharynx (Figure S2B-C). However, we noticed that in all cases, aberrant *npp-1* cells appeared in the brains of regenerating animals (Figure 1B-C; Figure S2D). These *npp-1*-positive ‘ectopic pharynx neurons’ (EPNs) in head fragments were unexpected. While the primary challenge of head fragments is to regenerate posterior tissue, the brain does not undergo drastic morphological changes, raising the possibility that EPNs might have been present prior to amputation. Indeed, in homeostatic *roboA*(RNAi) animals, we detected EPNs in the head (Figure 1D), indicating that EPNs arise independently of injury. To determine if *roboA* knockdown alters other pharynx cell types, we examined markers of pharynx muscle and epithelial cells in homeostatic animals. We detected multiple pharynx neuron and muscle markers expressed with a similar bilateral, brain-specific distribution (Figure 1E). We only detected ectopic pharynx epithelial cells in ectopic pharynges (Figure S1E-F), but not in the brain (Figure 1F). Furthermore, ectopic pharynges never expressed the pouch marker *dd_1320*, which may explain why ectopic pharynges are nonfunctional ^21^ (Figure 1F, S1E-F). In the brain, EPNs and pharynx muscle markers did not overlap (Figure 1G), suggesting that these are distinct cells, rather than cells with a mixed fate. Together, these results indicate that *roboA* restricts the maturation of multiple pharynx cell types to the proper anatomical region.

### EPNs in the head arise from stem cells, not mispatterned neurons

Robo receptors are best known for their roles in nervous system patterning ^28^. Because regeneration of a new head requires significant neurogenesis and nervous system remodeling, one possibility is that the EPNs appearing in the brain arise downstream of abnormal patterning ^21^. To separate the appearance of EPNs from neural patterning defects, we immunostaining of the optic nerves with Arrestin as a proxy for brain morphology. Importantly, uninjured, homeostatic *roboA*(RNAi) animals had normally patterned optic nerves (Figure 1H-I). As expected, the optic nerve in regenerating *roboA*(RNAi) animals displayed defasciculation and increased numbers of photoreceptors as compared to controls. To evaluate the impact of our knockdown, we performed quantitative RT-PCR and verified that knockdown animals had decreased *roboA* transcript (Figure S2G).Therefore, we conclude that *roboA* normally prevents the formation of EPNs independently of a mispatterned nervous system.

By *in situ* hybridization, *roboA* transcript appears enriched in the nervous system, but single-cell sequencing revealed a ubiquitous, low-level expression across all major cell types, including stem cells (Figure S3A-B). Between the brain lobes in wildtype animals, we identified *piwi-1^+^*stem cells that co-expressed *roboA*, indicating that *roboA* is present in stem cells (Figure S3C). To determine whether the appearance of EPNs in *roboA*(RNAi) animals requires stem cells, we exposed animals to ionizing radiation, which eliminates all stem cells ^29,30^. Then we knocked down *roboA* and challenged head fragments to regenerate a new pharynx (Figure S3D). Three days later, we failed to detect EPNs in the heads of irradiated *roboA*(RNAi) animals (Figure S3E-F), while they did appear in unirradiated controls. Therefore, the formation of EPNs requires active stem cells.

### *foxA* is required for pharynx neuron and epithelium but not muscle fates

The pioneer transcription factor *foxA* is required for pharynx regeneration through its expression in a subset of stem cells ^17^. *foxA^+^* stem cells are excluded from the brain and tail (Figure 1A) ^26^. Surprisingly, in the heads of *roboA*(RNAi) animals, we detected cells expressing both *foxA* and *npp-1* among the EPNs (Figure 2A). All *npp-1*-positive EPNs co-expressed *foxA* (59 cells from 3 animals), indicating that *roboA* normally suppresses *foxA* expression in the brain. To determine whether *foxA* is required for the appearance of EPNs, we knocked down *roboA* and *foxA* simultaneously. As compared to *roboA* knockdown alone, double knockdown animals had significantly reduced numbers of EPNs (Figure 2B-C). By contrast, the number of *laminin^+^* muscle cells significantly increased in double knockdown animals, indicating that ectopic pharynx muscle cells originate from stem cells independently of *foxA*. Together, these results suggest that *roboA* restricts stem cells from adopting a pharynx neuron fate in the brain by suppressing *foxA* expression.

**Figure 2.**
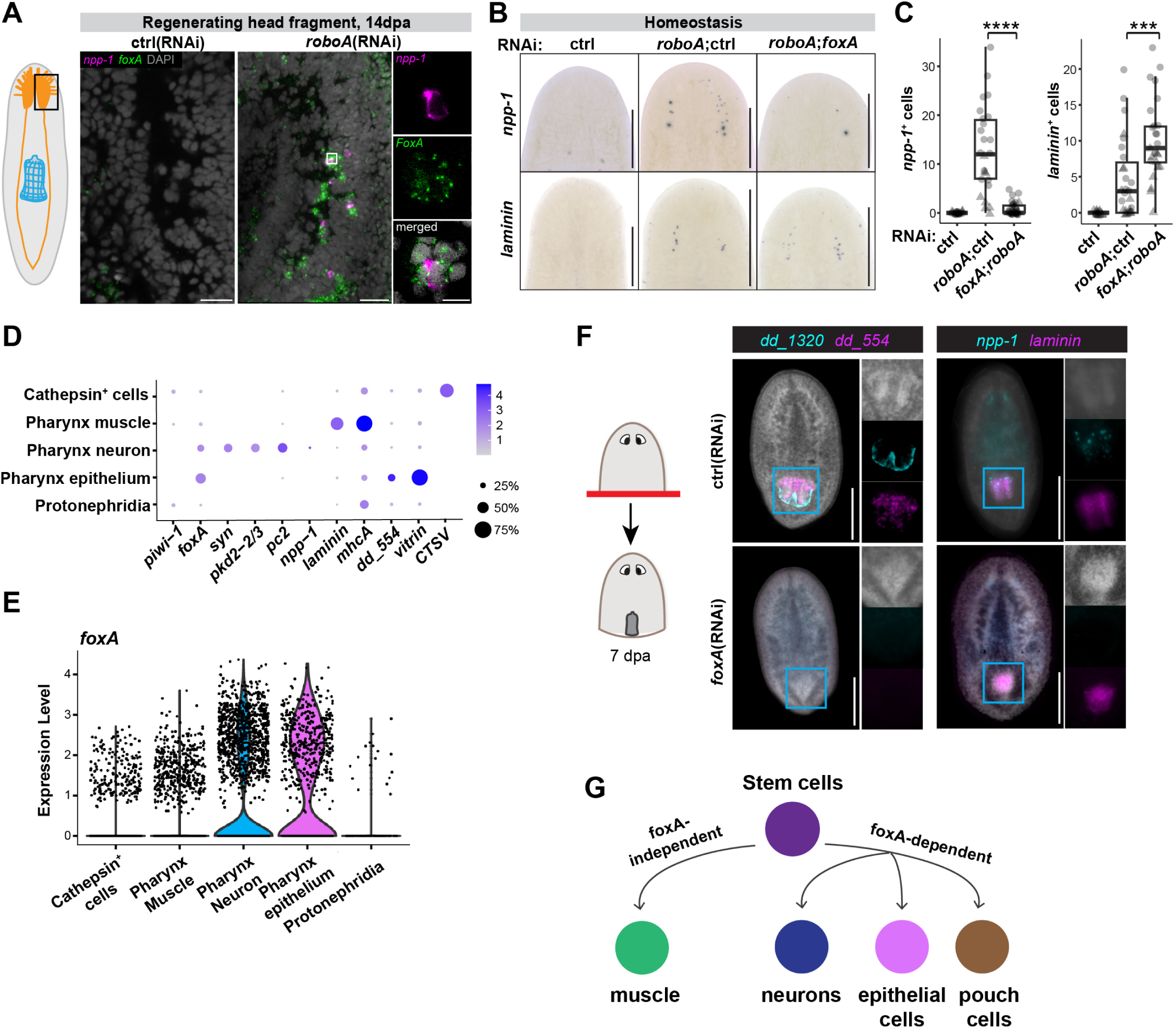
*foxA* is required for most pharynx fates. (A) Double fluorescent *in situ* hybridization (FISH) for *npp-1* (magenta) and *foxA* (green) in a newly regenerating head, 14dpa, with RNAi treatments as indicated. The white box highlights a single EPN expressing *foxA* and surrounded by ectopic *foxA*^+^ cells, with zoomed images on the right. DAPI (gray) stains nuclei. Scale bar = 100 μm; zoomed image = 20 μm. (B) ISH for *npp-1* (top row) and *laminin* (bottom row) in homeostatic RNAi animals as indicated. Scale bar = 200 μm. (C) Quantification of (B). ***, p≤0.001;****, p≤0.0001; one-way ANOVA with Tukey test. (D) Dot plot showing expression levels of pharynx cell type markers across different annotated clusters in the pharynx library (Figure S1C). Expression levels are indicated by color intensity, and dot size represents the percentage of cells expressing the marker. (E) Violin plot showing *foxA* expression levels across different annotated clusters in the pharynx library, with the highest expression in neurons and epithelial cells. (F) Regenerating head fragments in RNAi animals as indicated, 7 dpa, stained for pharynx pouch (*dd_1320*, cyan) and epithelium (*dd_554*, magenta); or pharynx neuron (*npp-1*, cyan) and muscle (*laminin*, magenta). *fox-A*(RNAi) animals fail to express *dd_554* and *dd_1320* (n=12, 100%); *foxA*(RNAi) animals fail to express *npp-1* but retain *laminin* (n=5, 100%). All control animals express these markers (n≥5, 100%). Scale bar = 200 μm. (G) Model of *foxA*-dependent and *foxA*-independent cell differentiation in the pharynx.

The differential requirement of ectopic neurons and muscles in *roboA*(RNAi) animals prompted us to examine the necessity of *foxA* for pharynx cell types during normal pharynx regeneration. In single-cell RNAseq analysis of isolated pharynx libraries, *foxA* was primarily enriched in pharynx neurons and epithelium (Figure 2D-E). To determine what cell fates require *foxA* during regeneration, we examined pharynx-specific markers in regenerating head fragments, which must form a new pharynx *de novo*. Seven days post-amputation, *foxA*(RNAi) animals completely lacked neuron (*npp-1*) and epithelial (*dd_554* and *dd_1320*) markers but retained muscle cells (*laminin*) in the position where the pharynx should be forming (Figure 2F). Together, we conclude that *foxA* is necessary for the production of pharynx neurons and epithelial cells, but not muscle cells (Figure 2G). Moreover, our results suggest that pharynx neurons and muscles might arise from different stem cell subtypes, which are both suppressed by *roboA* in the brain.

### Anterior-posterior patterning occurs independently of *roboA*

Ectopic expression of *foxA* in the brain could arise from a global disruption in axial patterning. In planarians, axial patterning is controlled by position control genes (PCGs), which are expressed in defined anatomical positions across the animal ^7^. Certain PCGs (*ndl-3*, *ptk-7*, and *wnt-p2*) regulate body axis patterning and determine pharynx position. Their knockdown shifts the body axis posteriorly, expanding the domain of *foxA* expression and promoting ectopic pharynx growth posterior to the normal pharynx ^8,10^. To determine whether global patterning is altered by *roboA* knockdown, we examined the expression of *sfrp-1*, *ndl-3*, *ptk-7*, and *wnt-p2* in homeostatic *roboA*(RNAi) animals. We detected no differences in their expression patterns (Figure S4). This indicates that the formation of EPNs in *roboA*(RNAi) animals are not due to defects in overall body patterning but instead arise from altered stem cell plasticity in the brain.

### *anos1* regulates pharynx neuron fates in concert with *roboA*

Our results suggest that RoboA, a transmembrane receptor, might act on stem cells to receive extracellular cues from the homeostatic brain. The canonical ligands of Robo are Slit proteins ^28,31,32^. To investigate whether RoboA functions via Slit, we knocked down the sole *slit* gene in planarians, and stained for *npp-1*. Surprisingly, *slit*(RNAi) animals exhibited no EPNs in the brain, suggesting that *roboA* regulates stem cell differentiation in a *slit*-independent manner (Figure 3A).

**Figure 3.**
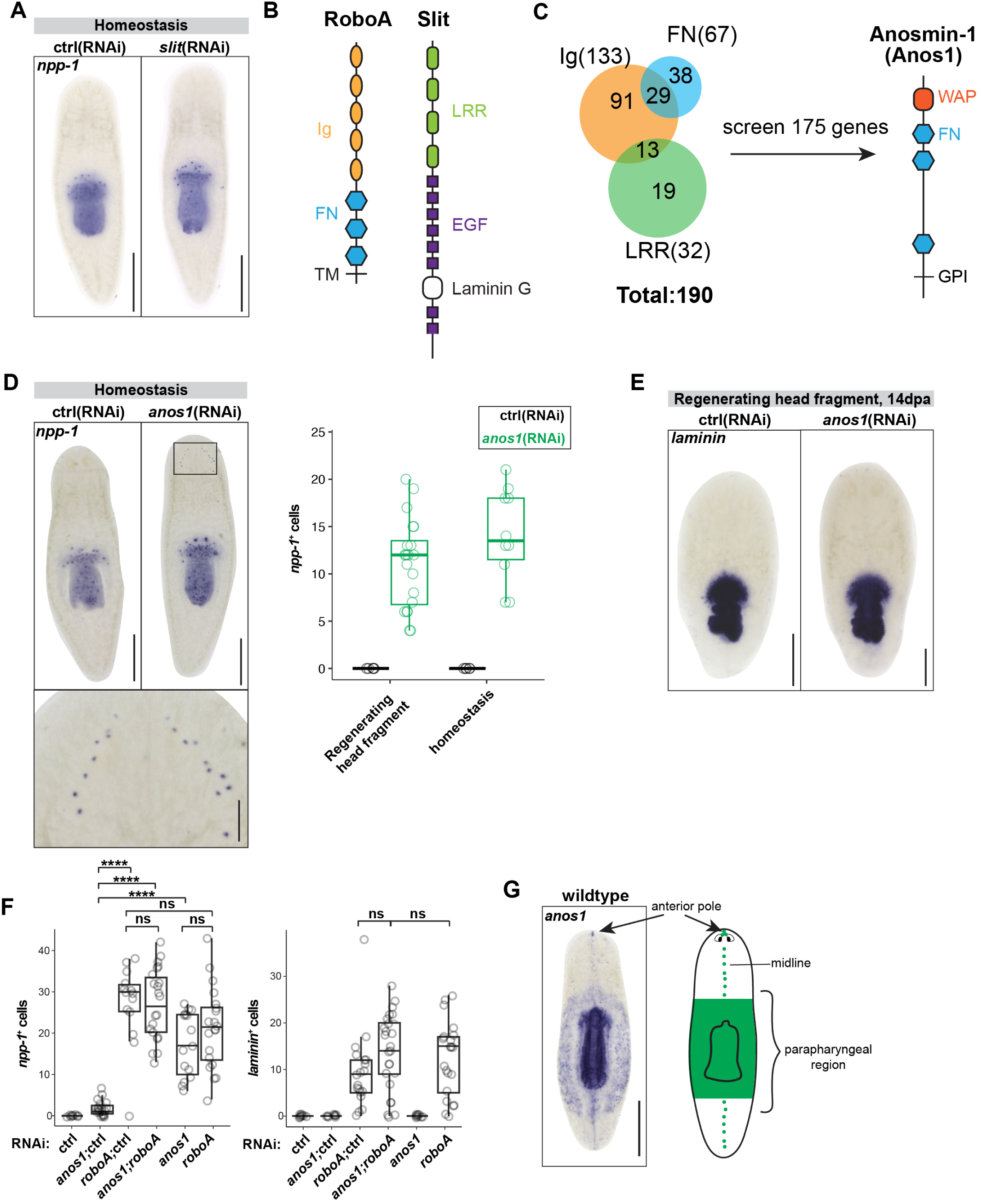
*anos1* knockdown phenocopies RoboA knockdown. (A) ISH of *npp-1* in homeostatic RNAi animals treated as indicated. No EPNs were detected (n≥15 animals). Scale bar = 500 μm. (B) Domain structure of RoboA and Slit proteins. RoboA contains immunoglobulin (Ig), fibronectin (FN), and transmembrane (TM) domains. Slit contains leucine-rich repeats (LRR), epidermal growth factor (EGF), and laminin G domains. (C) Domain-based screening for RoboA ligands included 175 candidate proteins with Ig, FN, or LRR domains. Right, Anosmin-1 (Anos1) protein contains a whey acidic protein domain (WAP), 3 FN domains and a GPI anchor. (D) Left, ISH of *npp-1* showing EPNs in homeostatic RNAi animals (zoomed in below). Scale bar = 500 μm (top), 100 μm (bottom). Right, quantification of EPNs in regenerating head fragments and homeostatic animals treated with control or *anos1*(RNAi). (E) ISH of *laminin* in regenerating head fragments from RNAi animals with no ectopic muscle cells. Scale bar = 500 μm (top). (F) Quantification of EPNs and ectopic muscle cells in homeostatic animals treated with RNAi as indicated. ns = not significant; ****p<0.0001; one-way ANOVA with Tukey test. Statistical comparisons of the control and other groups were omitted due to zero variance. Knockdown of *anos1* shows a dose-dependent effect in numbers of EPNs, where the *anos1*(RNAi) group has significantly more EPNs than *anos1*;ctrl(RNAi). By contrast, knockdown of *roboA* does not show a dose-dependent effect in numbers of EPNs or ectopic muscle cells. (G) ISH of *anos1* in wildtype animals showing expression in the parapharyngeal region, midline, and anterior pole, as shown in the schematic on the right. Scale bar = 500 μm.

RoboA is composed of five extracellular immunoglobulin domains, three fibronectin domains, and a transmembrane domain (Figure 3B). Robo receptors interact with the leucine-rich repeats of Slit proteins, mediating guidance and differentiation signals ^33^. Thus, to identify alternative ligands for RoboA, we searched the *S. mediterranea* genome for proteins with a signal sequence or transmembrane domain along with immunoglobulin, fibronectin, or leucine-rich repeat domains, finding 190 gene candidates (Figure 3C) (Supplementary Table 1). We screened 175 of these genes with RNAi, induced regeneration, and assessed animals 14 days later using ISH for EPNs. Knockdown of one gene, *anosmin-1* (*anos1*), phenocopied *roboA*(RNAi) (Figure 3D-E, Figure S5A-B) by inducing supernumerary pharynges and EPNs in both regenerating and homeostatic animals (Figure S5C-E). Unlike *roboA*(RNAi) animals, however, *anos1*(RNAi) animals did not express ectopic pharynx muscle (Figure 3E-F). This suggests that *anos1* specifically regulates neuronal differentiation in the brain.

Anos1 is a secreted protein containing a whey acidic protein (WAP) domain, three fibronectin domains and a GPI anchor (Figure 3C). Notably, mutations in human *ANOS1* (also referred to as *KAL-1*) are responsible for X-linked Kallmann Syndrome, a disorder characterized by hypogonadotropic hypogonadism and anosmia ^34,35^. Planarians have four ANOS1 homologs (Figure S5F), one of which has been previously characterized as a marker of ventral epithelial cells ^36,37^. The other three ANOS1 homologs are expressed in the brain (Figure S5G). These were included in the screen but failed to phenocopy *roboA* knockdown.

To determine whether *roboA* and *anos1* act in parallel or in the same pathway, we knocked both genes down simultaneously. We expected that if they act in parallel, we would detect increased numbers of EPNs as compared to single knockdowns (*roboA*;ctrl or *anos1*;ctrl). Double knockdown of *anos1* and *roboA* (*anos1*;*roboA*) did not lead to a significant increase in the number of EPNs compared to the *roboA*;ctrl group, but is significantly increased when compared to the *anos1*;ctrl group (Figure 3F). We observed a similar trend with *laminin*^+^ muscle cells when comparing *anos1*;*roboA* to *roboA*;ctrl, except that we detected no *laminin*^+^ cells in *anos1* single knockdown animals (Figure 3F). Therefore, these results suggest that RoboA and Anos1 likely function within the same pathway, potentially as a ligand-receptor pair. However, because Anos1 does not appear to be required for inducing pharynx muscle cells, it may also act in a parallel pathway.

### *roboA*-*anos1* signaling acts locally in the brain

To determine where *anos1* might act in the animal, we examined *anos1* transcript distribution. We found that it was expressed in the anterior pole, midline, pharynx muscle, and parapharyngeal regions (Figure 3G). The parapharyngeal distribution roughly mirrors *foxA* expression in this region, where *foxA*^+^ stem cells are located (Figure 1A). This observation raised the possibility that EPNs in *roboA* or *anos1* knockdown animals might arise from *foxA*^+^ stem cells that migrate into the brain and differentiate abnormally. Planarian stem cells have been shown to migrate over short and long distances ^38–40^, raising the possibility that parapharyngeal *anos1*^+^ normally suppresses this long-distance migration of *foxA^+^* stem cells.

To test this hypothesis, we surgically separated parapharyngeal tissues by head amputation and further prevented their regeneration through β*-catenin* knockdown (Figure 4A). In this background, the entire animal is anteriorized by an overall reduction in Wnt signaling, resulting in a loss of parapharyngeal identity ^9,41,42^. We reasoned that if EPNs originate from migrating stem cells, they would not appear in β*-catenin* knockdown head fragments lacking parapharyngeal cells. Alternatively, if they arise from local stem cell differentiation, EPNs would still appear.

**Figure 4.**
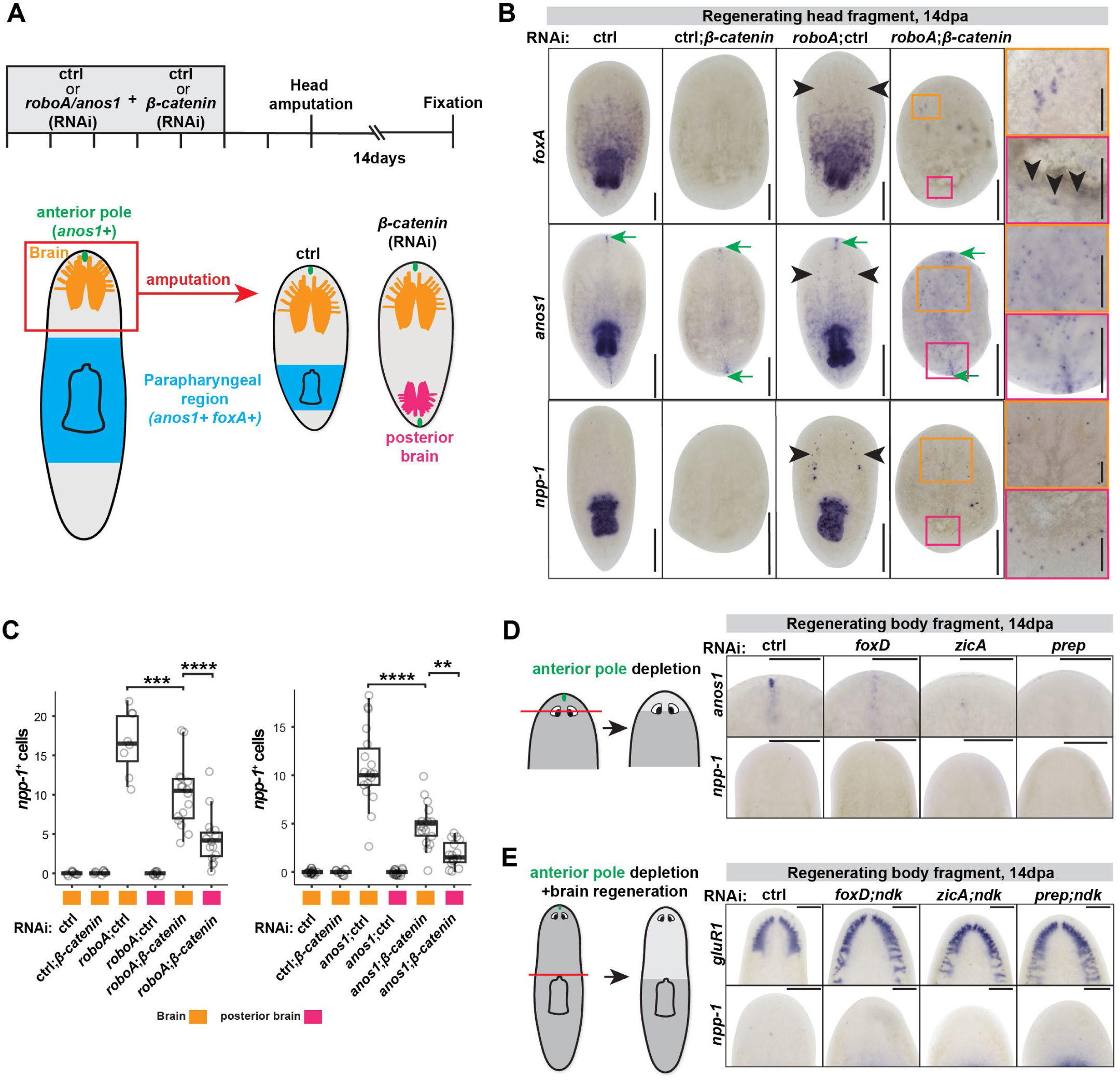
RoboA and Anos1 act locally in the brain. (A) Schematic of experimental design. Feeding of either control, *roboA*, or *anos1* RNAi preceded the administration of control or β-catenin RNAi. Two days later, animals were decapitated and fixed 14 days later. (B) ISH of *foxA* (top), *anos1* (middle), and *npp-1* (bottom) in head fragments treated with RNAi as indicated. *β-catenin*(RNAi) induces brain and anterior pole formation (green arrows) in the posterior, which persists in double knockdowns with *roboA*. Magnified views show clusters of ectopic cells (black arrowheads) in either the anterior (orange) or posterior (pink) brains. Scale bar = 250 μm, magnified views = 100 μm. (C) Quantification of EPNs from (A) treated with RNAi as indicated. **, p≤0.01; ***, p≤0.001;****, p≤0.0001; one-way ANOVA with Tukey test. (D) ISH of *anos1* and *npp-1* in RNAi animals treated as indicated to deplete the anterior pole. Scale bar = 250 μm. (E) ISH of *gluR1* and *npp-1* in RNAi animals treated as indicated to deplete the anterior pole and maximize brain size with *nou-darake* (*ndk*) knockdown. Scale bar = 250 μm.

We confirmed that this approach suppressed parapharyngeal identity, as *foxA* and *anos1* expression were lost following β*-catenin* knockdown (Figure 4B). In the double *roboA;*β*-catenin* knockdown context, the number of EPNs was significantly reduced in the anterior head (Figure 4B-C), suggesting that either residual parapharyngeal stem cells contribute to the formation of EPNs, or that EPN formation requires a certain level of Wnt signaling. Interestingly, we still detected EPNs and *foxA*^+^ cells in posterior-facing brains, but significantly fewer than in the anterior, likely due to their smaller size. This trend was consistent in *anos1*;β*-catenin* double knockdown animals (Figure 4C). Together, these findings demonstrate that *roboA* and *anos1* can act locally within the head to restrict stem cell differentiation, and EPNs arise wherever the brain is.

### Depletion of the anterior pole is not sufficient to induce ectopic pharynx fates

Since *roboA* and *anos1* suppress pharynx fates within the head, another potential source of Anos1 is the anterior pole, a group of muscle cells required for establishing anterior identity during regeneration ^43,44^. To test whether the anterior pole is required to suppress EPNs in the brain, we depleted it with a combination of surgical amputation and knockdowns of key anterior pole transcription factors, *foxD, prep,* or *zicA* ^45–47^. To ensure the removal of residual anterior pole signal, we surgically removed tissue anterior to the photoreceptors. If the anterior pole provides the inhibitory signal, we expected to observe the appearance of EPNs without any knockdown of *roboA* or *anos1*. However, despite successful anterior pole depletion, as validated by lack of *anos1* expression, these knockdown animals did not develop EPNs (Figure 4D), indicating that Anos1 at the anterior pole is not the source of inhibition.

One possible explanation for the absence of EPNs following anterior pole depletion is that residual *anos1* expression in the remaining brain tissue is sufficient to suppress EPN formation. To address this possibility, we forced regeneration of the entire brain by amputating as close to the pharynx as possible. Because knockdown of *foxD*, *prep*, or *zicA* results in generally smaller brain size, this may limit the potential induction and detection of EPNs. To overcome this limitation, we performed double knockdown experiments using *nou-darake* (*ndk*), a fibroblast growth factor receptor-like gene. *ndk* knockdown expands brain size ^48^ and can partially rescue anterior defects caused by *prep* knockdown^46^. In double knockdown animals (*foxD*;*ndk*, *prep*;*ndk*, or *zicA*;*ndk*), even though expression of the brain marker *gluR1* was restored, we did not observe any EPNs (Figure 4E). We conclude that the inhibitory signal that limits formation of EPNs does not originate in the anterior pole, but rather comes from elsewhere in the brain.

### *foxA* is the molecular switch for neuron fates

Our data reveals a latent potential of stem cells in the brain, which normally adopt brain-specific fates, to instead adopt pharynx-specific fates. When RoboA signaling is removed, stem cells reveal this potential by unleashing *foxA* expression. We hypothesized that stem cells in other parts of the body may also exhibit this potential. Because stem cells in the pharynx region normally induce pharynx-specific fates via FoxA (Figure 5A), we tested whether inhibition of *foxA* could also reveal a latent potential of stem cells. To accomplish this, wee compared the transcriptional profiles of brain and pharynx neurons, and further characterized how *foxA* knockdown could impact neuron fates.

**Figure 5.**
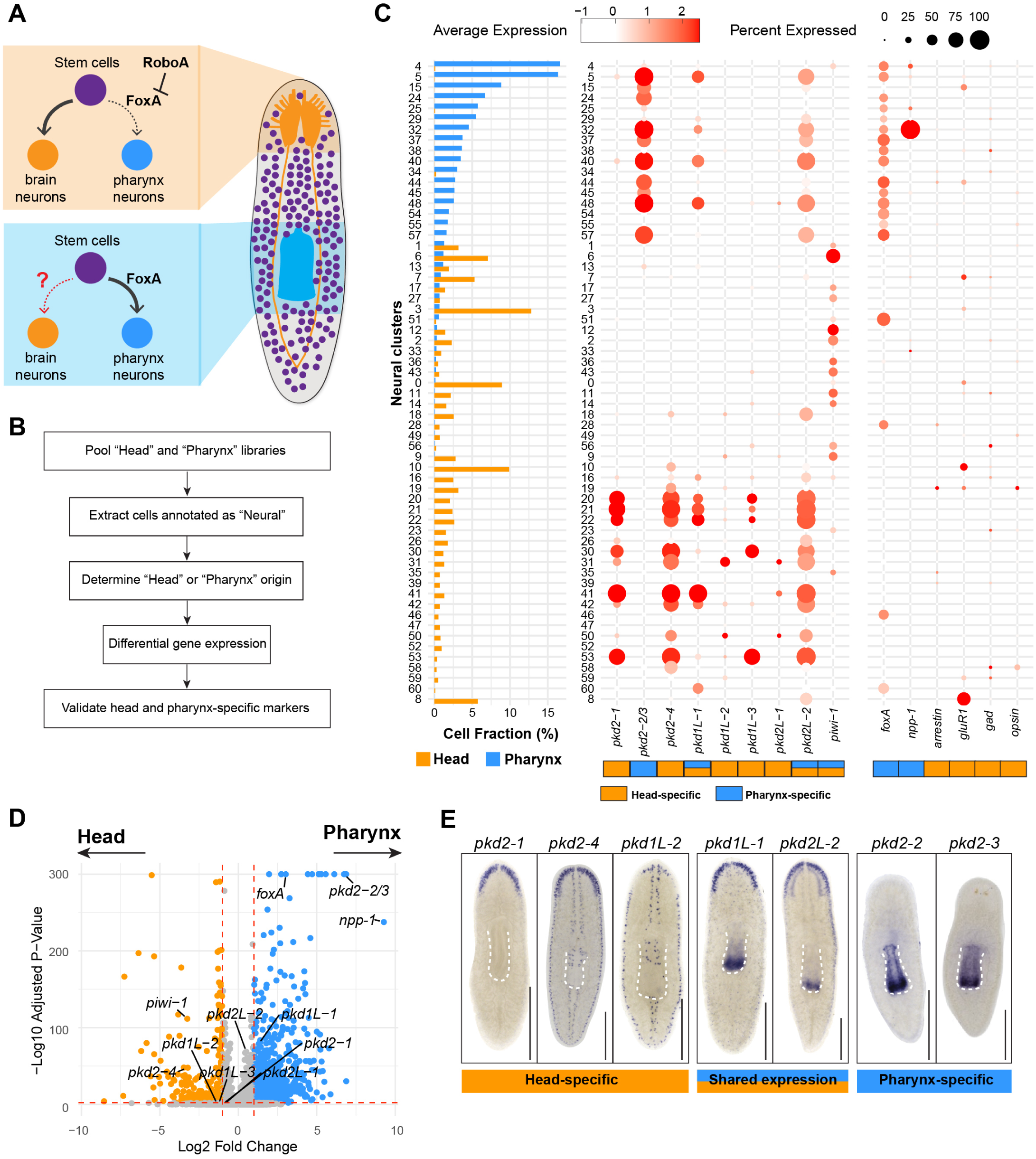
Transcriptomic distinction of pharynx and head neurons. (A) Schematic of regional differences in neuron differentiation from stem cells. (B) Single-cell RNA sequencing analysis workflow. (C) Left, bar plot showing the distribution of cells from either the head or pharynx library across neural clusters. Right, dot plot showing the expression of head-specific and pharynx-specific genes across neural clusters. Dot size represents the percentage of cells expressing each gene, and color intensity reflects expression levels. (D) Volcano plot showing differentially-expressed genes (DEGs) in head and pharynx neurons. Wilcoxon rank sum test, p-value adjusted by Bonferroni correction. Orange and blue dots represent DEGs with |log_2_ fold change| >1 and -log_10_(adjusted p-value)<0.01. (E) ISH of *pkd* family members in wildtype animals. Scale bar = 500 μm.

To identify unique molecular profiles for brain and pharynx neurons, we reanalyzed the published region-specific single-cell libraries ^49^, extracting neural-specific clusters (Figure 5B). Pharynx neurons and brain neurons formed distinct clusters ^49–51^. Of 61 clusters, 17 are primarily composed of pharynx neurons, while 24 consist only of head neurons (Figure 5C). As expected, pharynx neuron clusters had higher *foxA* expression (Figure 5C-D). Differential gene expression analysis using the Wilcoxon rank sum test identified members of the *polycystin* (pkd) gene family as unique to either the head or pharynx (Figure 5D). Consistent with a recent report and single-cell analysis, *pkd2-2* and *pkd2-3* show pharynx-specific expression, while other paralogues are predominantly expressed in auricles and brain regions (Figure 5E) ^51^. These findings suggest that brain and pharynx neurons are molecularly distinct, with *foxA* playing a key role in defining these cell states.

With these markers in hand, we analyzed the impact of long-term *foxA* depletion on differentiation of specific neuron populations. In this context, *foxA*(RNAi) animals lose their pharynges and instead develop a dorsal outgrowth (Figure 6A) enriched in neuronal markers such as *pc2* and *ndk* ^17^. We generated single-cell sequencing libraries from the parapharyngeal regions of *foxA*(RNAi) and control animals 21 days after RNAi administration (Figure 6B). Both libraries were pooled, clustered and annotated (Figure S6A-B). The primary difference between control and *foxA* knockdown animals was the absence of pharynx epithelium and pouch cells (Figure S6A-B), consistent with the loss of epithelial marker staining in animals examined 7 days post-amputation (Figure 2). Fractions of neurons in the control and *foxA*(RNAi) libraries were comparable, suggesting that *foxA*(RNAi) did not broadly impair neurogenesis (Figure S6B). Next, we examined neuron composition by calculating the fold change between the *foxA*(RNAi) cells relative to *ctrl*(RNAi) cells within each cluster (Figure 6C, left). We found that clusters enriched in *foxA*(RNAi) cells had higher expression of head-specific marker genes *pkd2-1* and *pkd2-4*, while clusters enriched in controls expressed high *foxA* but lacked head-specific markers, as expected (Figure 6C, right). These findings suggest that *foxA* knockdown induces a broad transcriptional shift toward a brain-like profile, indicating that *foxA* functions as a central regulator of differentiation, by integrating global inputs in stem cells.

**Figure 6.**
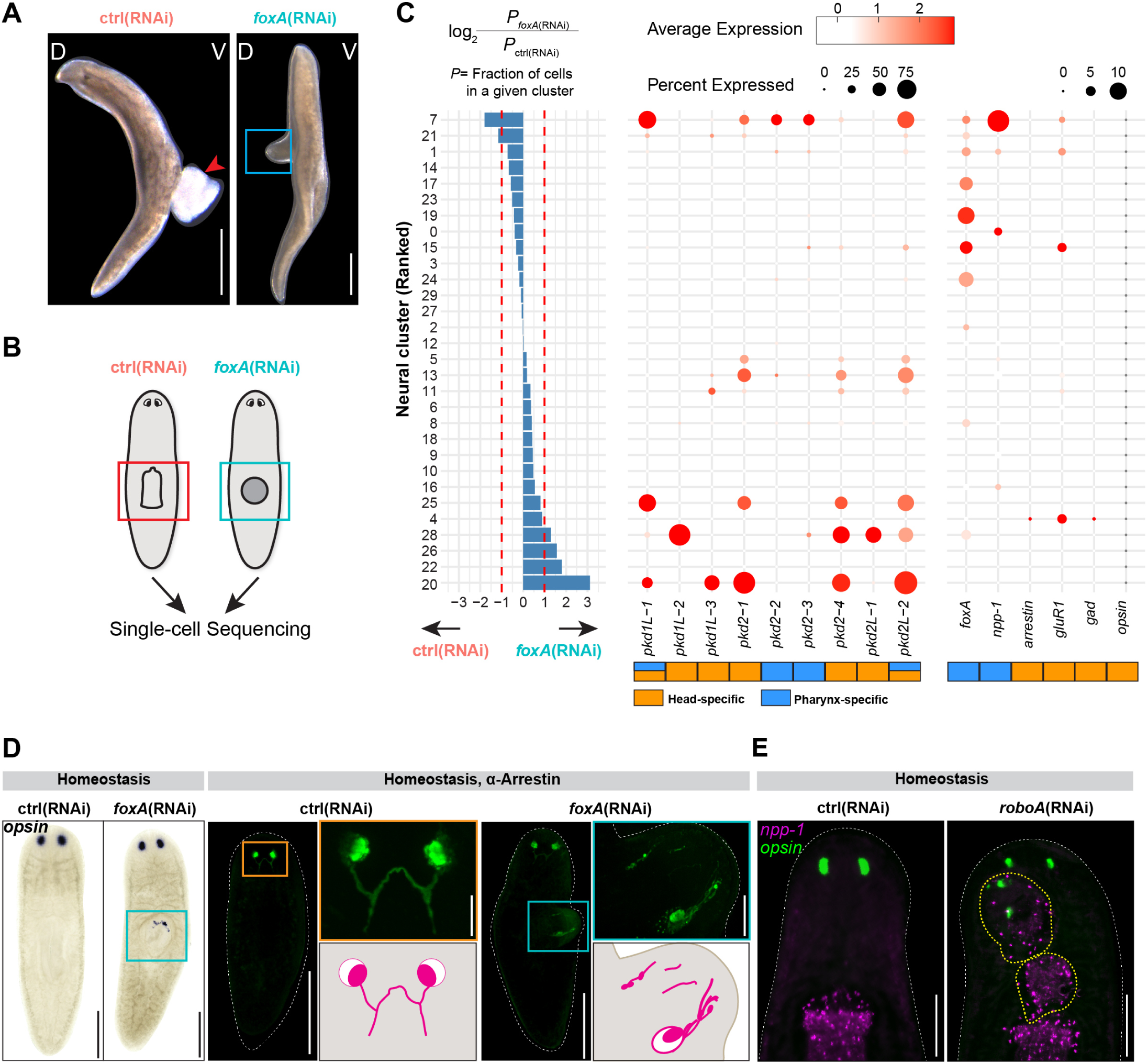
*foxA* dictates a fate switch between pharynx and brain neurons. (A) Live animals, 21 days after RNAi feeding. Animals were soaked in tricaine, which causes the pharynx to emerge (red arrowhead). *foxA*(RNAi) animals lack a pharynx and instead form a dorsal outgrowth (blue box). D = dorsal; V = ventral. Scale bar = 500 μm. (B) Schematic of tissue extracted from parapharyngeal region for single-cell RNA sequencing. (C) Left, log_2_ fold change in neural composition across clusters, comparing *foxA* over control(RNAi) animals. Right, Dot plot showing the expression of known marker genes across clusters. Dot size represents the percentage of cells expressing the gene, and color intensity reflects the expression level. (D) ISH of *opsin* (left) and immunostaining of Arrestin (right) in homeostatic animals treated with control or *foxA*(RNAi). Ectopic photoreceptor expression appears in the pharynx region (blue boxes) in *foxA*(RNAi) animals (50% *opsin*^+^, n=6; 28.6% Arrestin^+^, n=7; none of the control animals, n≥10). Insets show magnified views. Scale bar = 250 μm. Inset = 100 μm. (E) Double FISH for *npp-1* (magenta) and *opsin* (green) in homeostatic animals. The ectopic pharynx of *roboA*(RNAi) animals (yellow dashed line) contains both *opsin* and *npp-1* (10.5% *opsin*^+^ *npp-1*^+^, n=19 animals; 0/20 control animals). Scale bar = 250 μm.

To validate these findings, we examined the expression of several brain-specific markers in *foxA*(RNAi) animals with a dorsal outgrowth. Notably, these dorsal outgrowths had optic nerves and structured eye cups, indicating abnormal respecification to head-specific cell fates in the pharynx region (Figure 6D). We also detected head-specific markers *pkd2-1* and *pkd2-4*, along with brain-specific *glutamic acid decarboxylase* (*gad*) and *gluR1* in the dorsal outgrowth, where they are otherwise never expressed (Figure S6C). These results suggest that *foxA* normally maintains pharynx fates and suppresses latent brain fates.

We speculated that the boundaries between pharynx and brain regions might be blurred in animals with ectopic pharynges, such as *roboA* knockdown animals. Analysis of the cellular composition of cephalic outgrowths showed that it contains both the eye marker *opsin* and EPNs (Figure 6E). This mixing of pharynx and brain fates in the prepharyngeal region indicates that stem cells exhibit a shared plasticity to adopt either brain or pharynx fates, with *roboA* and *foxA* acting as molecular switches that reinforce spatially appropriate cell fate decisions.

## Discussion

Planarian stem cells can generate all the animal’s cell types ^6^, suggesting that they exhibit extreme plasticity. This plasticity of adult stem cells has been lost in most animals, along with their ability to regenerate robustly. Thus, the mechanisms that enable stem cell plasticity, or how it might be regulated to coordinate faithful regeneration, remain unclear. We provide the first evidence of latent adult stem cell plasticity in planarians, which depends on a newly identified potential ligand-receptor pair to spatially constrain cell fates through a pioneer factor-driven transcriptional response (Figure 7). Our study bridges the gap between the global signals known to broadly pattern the organism and downstream transcriptional responses in stem cells that are influenced by their surroundings. This mechanism involves a previously unrecognized triad — Anos1, RoboA, and FoxA — that refines stem cell plasticity to ensure that differentiation occurs in the appropriate anatomical position.

**Figure 7.**
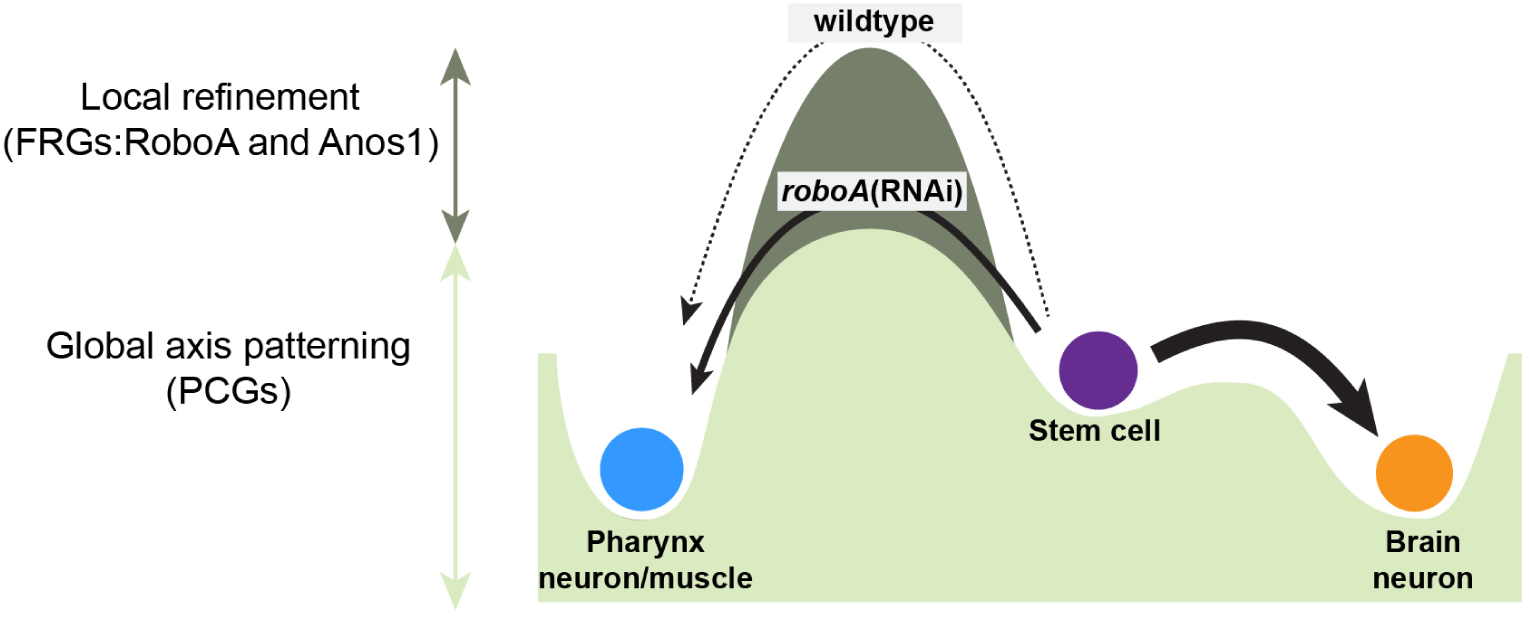
Model for stem cell plasticity modulated by RoboA and Anos1 in the brain. Extrinsic control of stem cell fate decisions in planarians includes two layers of regulation: global axis patterning by position control genes (PCGs) and local refinement by fate-reinforcing genes (FRGs) RoboA and Anos1. Following establishment of broad anatomical domains, RoboA and Anos1 function locally to suppress inappropriate differentiation, reinforcing the barrier preventing pharynx fate. In wildtype animals (dashed arrow), RoboA/Anos1 signaling prevents stem cells from adopting EPN and pharynx muscle fates in the brain. By contrast, *roboA*(RNAi) (solid arrow) lowers the barrier to stem cells, enabling the adoption of pharynx neuron and muscle fates in the brain. Notably, even in *roboA*(RNAi) animals, most stem cells still differentiate normally into brain neurons (thick arrow), highlighting the importance of integrating both PCGs and FRGs.

Local stem cell differentiation is controlled by position control genes (PCGs) and organ-maintenance signals (known as the target zone) ^1^. RoboA’s function aligns with the target zone concept by acting independently of PCGs to suppress pharyngeal fate in the brain, a downstream product of global axis signals. Because PCG expression and overall animal morphology remain intact in *roboA*(RNAi) animals, our findings refute the idea that *roboA* is a PCG as previously proposed ^1^. Instead, we define genes that suppress plasticity without broadly perturbing global patterning as ‘Fate-Reinforcing Genes’ (FRGs) (Figure 7). In this framework, PCGs expressed in muscles define broad body regions. Downstream of PCGs, FRGs suppress inappropriate cell types, thereby maintaining proper organ positioning and maintenance. Interestingly, even in the absence of RoboA, anterior stem cells preferentially generate brain-specific neurons rather than pharynx neurons, further demonstrating that FRGs serve as modulators of cell identity rather than position determinants. This fine-tuning role highlights a previously underappreciated level of spatial control in stem cell fate decisions.

The cellular mechanisms that determine when and how stem cell plasticity becomes restricted are poorly understood. Planarian stem cells are heterogeneous, with subpopulations defined by fate-specific transcription factors (FSTFs) ^18,52,53^. Recent work proposed that these subpopulations arise in a cell-cycle dependent manner ^54^. Stem cells in G0 are naive and lack FSTFs, while G2-phase cells express FSTFs and give rise to lineage-committed progenitors. These G2 cells are stochastically distributed across the animal body, suggesting that progenitors might migrate long distances to their final destination ^40^. By contrast, our findings support a model of locally-constrained plasticity, where regional signals limit the range of potential fates. Comparisons with amphibian regeneration offer insight into how different systems resolve plasticity. In axolotl limb and *Xenopus* tail regeneration, Schwann cells and connective tissue cells can dedifferentiate into progenitors, but these progenitors are intrinsically constrained in their potency to their own lineage ^2,55^. Our findings highlight a distinct strategy in planarians, where adult stem cells maintain broader plasticity and fate is instead refined by local extrinsic cues. We propose that in highly regenerative organisms, stem cell differentiation is not strictly pre-determined, but actively sculpted by the tissue environment.

Although Robo signaling is mediated through interaction with Slit ligands, our findings demonstrate that RoboA regulates stem cell differentiation in planarians independently of Slit. Molecular evidence from other systems has shown that Robo has a broader set of possible interactions beyond Slit. For example, Robo can interact with non-Slit proteins such as Netrin receptors and other Robo family members ^24,56,57^, homodimerize ^58^ or be cleaved to activate binding partners ^59^. Intriguingly, a recent interactome study suggests that the *C. elegans* Robo homolog Sax-3 can bind to multiple proteins *in vitro*, including Slit and Anos1 ^60^. Our results contribute Anos1 as a new alternative Robo ligand candidate, a new organism, and a novel cellular context to the known possible Robo functions.

ANOS1 mutations cause Kallmann syndrome, characterized by anosmia and hypogonadotropic hypogonadism, thought to result from impaired migration of olfactory and GnRH neurons ^61–63^. While the absence of Anos1 homologs in rat and mouse genomes complicates mechanistic studies, Anos1 modulates FGF signaling through interactions with FGF ligands and heparan sulfate, as observed in zebrafish ^64^ and roundworms ^65–68^. A Robo mutation found in a Kallmann syndrome patient ^69^, combined with our findings identifying Anos1 as a potential RoboA ligand, suggests a previously unrecognized and conserved role for Robo-Anos1 interactions in neural differentiation, highlighting new molecular insights relevant to Kallmann syndrome.

## Limitations of the study

Our model proposes that RoboA acts on stem cells, but we have only examined this with *in situ* hybridization, leaving the localization of RoboA and Anos1 elusive. Moreover, we cannot rule out whether de-differentiation or transdifferentiation accounts for the formation of EPNs, although there is no evidence for these types of fate transitions in planarians.

## Materials and methods

### Animal husbandry

*Schmidtea mediterranea* asexual clonal line CIW4 was maintained in a recirculating water system supplemented with Montjuïc salts ^70^. Animals were fed beef liver and cleaned weekly. Before use, they were transferred to a static culture with 50 µg/mL gentamicin, and starved for at least one week.

### RNA interference

RNA interference was conducted by feeding animals with either bacterial-expressed or in vitro synthesized double stranded RNA. For bacterial RNAi, cDNA of the target genes was cloned into either the pJC53.2 or T4P vectors and transformed into HT115 competent cells ^71^. Cultures were grown in 2XYT broth at 37°C with shaking at 200–250 rpm until reaching an OD600 of 0.6, followed by induction with 1 mM IPTG and additional shaking for 2 hours. Bacterial cultures were then pelleted, and every 50 mL culture pellet was resuspended in 125 µL of liver paste before being administered to the animals. For double RNAi, equal volumes of two bacterial cultures were mixed before pelleting. Bacterial RNAi was given every other day for a total of four feedings for single gene knockdown and eight feeds for double gene knockdown unless otherwise indicated in the figure. All primer sequences are listed in Supplementary Table 1.

Screening of roboA ligands utilized *in vitro-*synthesized dsRNA. Most dsRNA was generated using T7-overhang primers (Supplementary Table 2) to amplify candidate genes from cDNA, and the PCR products were used as templates for in vitro transcription ^72^. The dsRNA was then mixed into a 4:1 liver-to-water paste, with 4 µg of dsRNA per 10 µL of liver. Exceptions to these in the screen were *slit*, *roboA*, *roboB*, *roboC* and *roboD*, which were cloned into pJC or T4P vectors and administered with bacterial feeding. Animals were fed every other day for a total of six feedings. Animals were either amputated 5 days after the final RNAi feeding and fixed 14 days later, or maintained without injury for 14 days, unless otherwise indicated in the figures. All experiments used the *Caenorhabditis elegans* gene *unc-22* as a control.

For stem cell depletion, planarians were exposed to 60 Gray using a J.L. Shepherd & Associates Mark I-68 Irradiator. Following exposure, animals were rinsed immediately, transferred to fresh water, and fed RNAi food.

### Genome-wide identification of IgSF, FN, and LRR domain containing proteins

Sequences derived from the dd-Smed-v6 transcriptome were utilized for gene annotation ^73^. Initially, transcripts, including all isoforms, were translated into protein sequences and subsequently filtered to retain only those sequences with a minimum length of 100 amino acids. Conserved domain annotations were conducted using the SUPERFAMILY database through InterProScan (version 5.71) ^74^. Proteins exhibiting immunoglobulin, fibronectin, or leucine-rich repeat domains underwent further analysis for the presence of signal peptides and transmembrane domains using DeepTMHMM (1.0.4) ^75^. 196 proteins from 190 genes met these criteria, and the longest isoforms of each gene were further used for primer design. PCR failed for 15 of these genes.

### qRT-PCR

For each biological replicate, 5 animals were homogenized in Trizol (Thermo Fisher 15596018) using Lysing Matrix D Tubes (MP Biomedicals 116913100) and a Bead Bug homogenizer (Benchmark). RNA was extracted according to the standard Trizol protocol. cDNA was synthesized using Superscript VILO (Life Technologies 11754250). PCR reactions were prepared with SYBR Green and GAPDH was used as a control. Reactions were performed on an Applied Biosystems Viia7 Real-Time PCR System. Each biological replicate included three technical replicates, and Ct values were averaged. Data were analyzed using the Delta-Delta Ct method. All primers can be found in Supplementary Table 1.

### *In situ* hybridization and immunostaining

*In situ* hybridizations were conducted as previously described with some modifications ^76,77^. Fixed animals were rehydrated, bleached (5% formamide, 6% H2O2, 0.5% SDS in 0.5× SSC) for 1 hour, and treated with proteinase K [4 μg/ml in PBS with 0.3% Triton X-100 (PBSTx), Thermo Fisher 25530049] for 10 minutes, followed by a 10-minute fixation in 4% formaldehyde. Probes were added and incubated at 56°C for 16 hours following 2 hours of pre-hybridization. Samples were washed twice in wash hybe (20 minutes each), followed by 1:1 wash hybe:2× SSC-0.1% Tween 20 (20 minutes), 2× SSC-0.1% Tween 20 (30 minutes), and 0.2× SSC-0.1% Tween 20 (30 minutes) at 60°C, and then washed three times for 10 minutes each with MABT (100mM maleic acid, 150mM NaCl, 0.1% Tween-20, pH to 7.5 with NaOH) at room temperature. The animals were blocked in a solution of 0.5% Roche Western Blocking Reagent and 5% inactivated horse serum in MABT for 2 hours at room temperature and incubated with antibody solution overnight at 4°C. Antibody solution was made with either 1:3000 anti-DIG-AP (Roche 11093274910), 1:1000 anti-DIG-POD (Roche 11207733910), or 1:1000 anti-FITC-POD (Roche 11426346910) diluted in blocking solution. Tyramide conjugates were synthesized following the method outlined in ^78^. Tyramide signal amplification was achieved by incubating planarians for 5–10 minutes in a 1:1000 dilution of fluorophore-conjugated tyramide in 100 mM borate buffer (pH 8.5, 2 M NaCl, 0.003% H_2_O_2_, and 0.1M Boric acid). For double FISH experiments, residual peroxidase activity was quenched by incubating the samples for at least 2 hours in 1% sodium azide in PBSTx ^77^. After development, animals were soaked and mounted in ScaleA2 (4 M urea, 20% glycerol, 0.1% Triton X-100, 2.5% DABCO). Some hybridizations were performed using a CEM InSituPro Hybridization robot up to the development stage, using the same protocol.

*In situ* hybridization chain reaction (HCR) was performed following the manufacturer’s mouse embryo protocol (Molecular Instruments) with slight modifications ^79^. Animals were fixed, bleached, and treated with proteinase K as described above. Samples were pre-hybridized in HCR™ Probe Hybridization Buffer (v3.0) at 37°C for 30 min, followed by overnight probe incubation at 37°C with a 16 nM probe solution in HCR™ Probe Hybridization Buffer (v3.0). Probes were designed using insitu_probe_generator ^80^ and ordered from IDT (Supplementary Table 3). Samples were then washed four times at 37°C for 15 min in HCR™ Probe Wash Buffer (v3.0), followed by two washes at room temperature for 5 min in 5× SSC with 0.1% Tween 20. Next, samples were incubated in HCR™ Amplifier Buffer (v3.0) at room temperature for 30 min. Fluorophore-conjugated hairpins (h1 and h2; 10 μL of a 3 μM stock each) were “snapped cool” by heating at 95°C for 90 sec and then cooled to room temperature in a PCR machine. The cooled hairpins were added to 500 μL of HCR™ Amplifier Buffer (v3.0) to prepare the hairpin solution, and samples were incubated with this solution in the dark at room temperature overnight. The following day, samples were washed four times in 5× SSC with 0.1% Tween 20 and mounted in 80% glycerol.

For immunostaining, animals were fixed and bleached as for *in situ* hybridizations. 1% BSA was used for blocking and antibody solution. A mouse anti-Arrestin antibody was used at a 1:1000 dilution ^81^, and an anti-mouse-Alexa conjugated antibody was used at a 1:2000 dilution.

### Cell sorting and 10X single-cell library preparation

To generate single-cell libraries, 50 trunks from control(RNAi) or *foxA*(RNAi) animals were collected 21 days post-feeding. The tissues were dissociated into single-cell suspensions by mincing in CMFB buffer [calcium-magnesium-free solution with 1% BSA (400 mg/L NaH2PO4, 800 mg/L NaCl, 1200 mg/L KCl, 800 mg/L NaHCO3, 240 mg/L glucose, 1% BSA, 15 mM HEPES, pH 7.3)] and gently rotating for 2 hours at room temperature. The cells were then centrifuged at 500 g for 5 minutes, resuspended, and passed through a 30 µm cell strainer (BD Biosciences, 340628) to remove clumps. The concentration of filtered cells was determined using a TC20 automated cell counter (Bio-Rad). After centrifugation, the cell concentration was adjusted to 100,000 cells/mL in a staining buffer [CMFB containing 0.4 µM Calcein-AM Cat#C3100MP] and incubated with gentle rotation at room temperature for 20 minutes. Live cells were identified by Calcein-AM staining and sorted using a Sony MA900 Cell Sorter. A total of 100,000 cells were sorted and diluted to 1000 cells/µL for subsequent 10X library preparation. To aim for recovery of 10,000 cells after sequencing, 16,500 cells were loaded onto the 10X Genomics Chromium Controller for subsequent library preparation using Chromium Next GEM Single Cell 3L Reagent Kit v3.1.

To deplete 16S sequence contamination from 10X libraries, Depletion of Abundant Sequences by Hybridization (DASH) method was used as previously described ^82^. Briefly, the cDNA library was treated with CRISPR/Cas9 targeting 16S at 37°C overnight. Following CRISPR treatment, the cDNA was purified and eluted into 15 µL using AMPure beads (Beckman) according to the manufacturer’s protocol. The cDNA was then diluted to 30 µL and re-amplified with cDNA primers (R1 + TSO) for 10 cycles. After PCR amplification, the cDNA was processed according to the 10X Genomics protocol for enzymatic fragmentation and indexing. Three biological replicates were included in this study. Libraries were pooled and sequenced on the Illumina NextSeq 2000 platform. To verify the removal of 16S cDNA, individual samples were analyzed using the Fragment Analyzer (Agilent).

### Single-cell analysis

Sequencing reads of single-cell libraries were aligned to a customized genome file containing both the chromosomal-level genome (Smed_chr_ref_v1)^83^ and 16S sequence (SMED30032887) by CellRanger (6.1.2) ^84^. Data preprocessing, including normalization and variable feature identification, was conducted using the Seurat (5.0.0) package ^85^ in R(4.1.1). Single-cell RNA-seq data matrices were read into Seurat objects using the Read10X function, and each dataset was normalized and processed to identify variable features.

The control(RNAi) and *foxA*(RNAi) datasets were combined into a single Seurat object and normalized using standard Seurat workflows. We used a previously published dataset ^49^ to assign cell type identities as a reference with Seurat’s FindTransferAnchors and TransferData functions. Before using it as a reference, the Fincher 2018 dataset’s gene annotations were converted from the dd-Smed-v4 transcriptome format to the SMESG gene model to match the datasets generated in this study ^73^. When multiple dd-Smed-v4 transcripts corresponded to the same SMESG gene, their read counts were aggregated. Cell type identities were predicted based on the “Major.cluster.description” annotations from the Fincher 2018 dataset, which define the main cell types. Cells predicted to be “Neural” were further extracted and subclustered by the first 20 Harmony coordinates used for clustering with the “FindClusters” function, set at a resolution of 2, employing the shared nearest neighbor (SNN) approach.

To assess the cell-type composition, cells were grouped based on their original sample type [control(RNAi) or *foxA*(RNAi)] and their predicted major cell type. Bar plots were then used to visualize and compare the relative abundance (log2 fold change) of each cell type between the control(RNAi) and *foxA*(RNAi) conditions, highlighting differences in cell-type fractions between the two experimental groups.

### Statistics and image quantification

Imaging was done using a Leica M165F with a DFC7000T camera for live animals, a Zeiss 710 confocal microscope, or a Keyence BZ-X800 microscope for colometic ISH and fluorescent stainings. ImageJ software (FIJI) was used to process images and for manual counting of cells. PRISM-Graphpad (10.4.0) was used to perform one-way ANOVA with Tukey test as indicated in figure legends, except in cases where the variance was zero, making the Tukey test inapplicable. *p<0.05; **p<0.01; ***p<0.001; and ****p<0.0001.

### Phylogenetic analysis

Sequences for phylogenetic analysis were retrieved from the NCBI database, with accession numbers detailed in Supplementary Table 4. Planarian sequences were specifically obtained from the dd-Smed-v6 transcriptome ^73^. The retrieved sequences were translated into proteins and aligned using CLUSTALW with the SLOW/ACCURATE setting. Phylogenetic trees were inferred using the Maximum Likelihood method ^86^ with bootstrap support (n=1000 replicates), implemented in IQ-TREE2 (version 2.2.6) ^87^. Final tree visualization and layout optimization were performed using FigTree (version 1.4.4).

## Resource availability

### Lead contact

For additional information or requests for reagents, please contact the lead contact, Carolyn Adler.

### Data and code availability

All data are available upon request. scRNALseq data have been deposited into NCBI Accession number GSE292456 (https://www.ncbi.nlm.nih.gov/geo/query/acc.cgi?acc=GSE292456). Codes have been deposited on Github (https://github.com/kw572/RoboA).

## Acknowledgments

This work was funded by a National Institutes of Health grant R01GM139933 (to C.E.A.), a Cornell University Stem Cell Program fellowship, Cornell University Center for Vertebrate Genomics Scholarship and Taiwan Ministry of Education scholarship (to K-T.W.). We thank the Cornell University Biotechnology Resource Center’s Flow Cytometry (RRID:SCR_021740), Imaging (RRID:SCR_021741), and Genomics (RRID:SCR_021727) cores; T. Inoue for Arrestin antibody; members of the Adler laboratory for insight; T. Tumbar for critical reading of the manuscript.

## Author contributions

K-T.W., Y-C.C. and C.E.A. conceived the study. C.E.A. supervised the research and acquired funding. K-T.W., Y-C.C., F-Y.T. and C.P.J. performed the ISH, RNAi experiments, and imaging. K-T.W. generated the sequencing data and performed all statistical analyses. K-T.W. and C.E.A. wrote the original draft. All authors edited, read, and approved the manuscript.

## Disclosure and competing interests statement

The authors declare that they have no conflict of interest.

---

**Figure S1:**
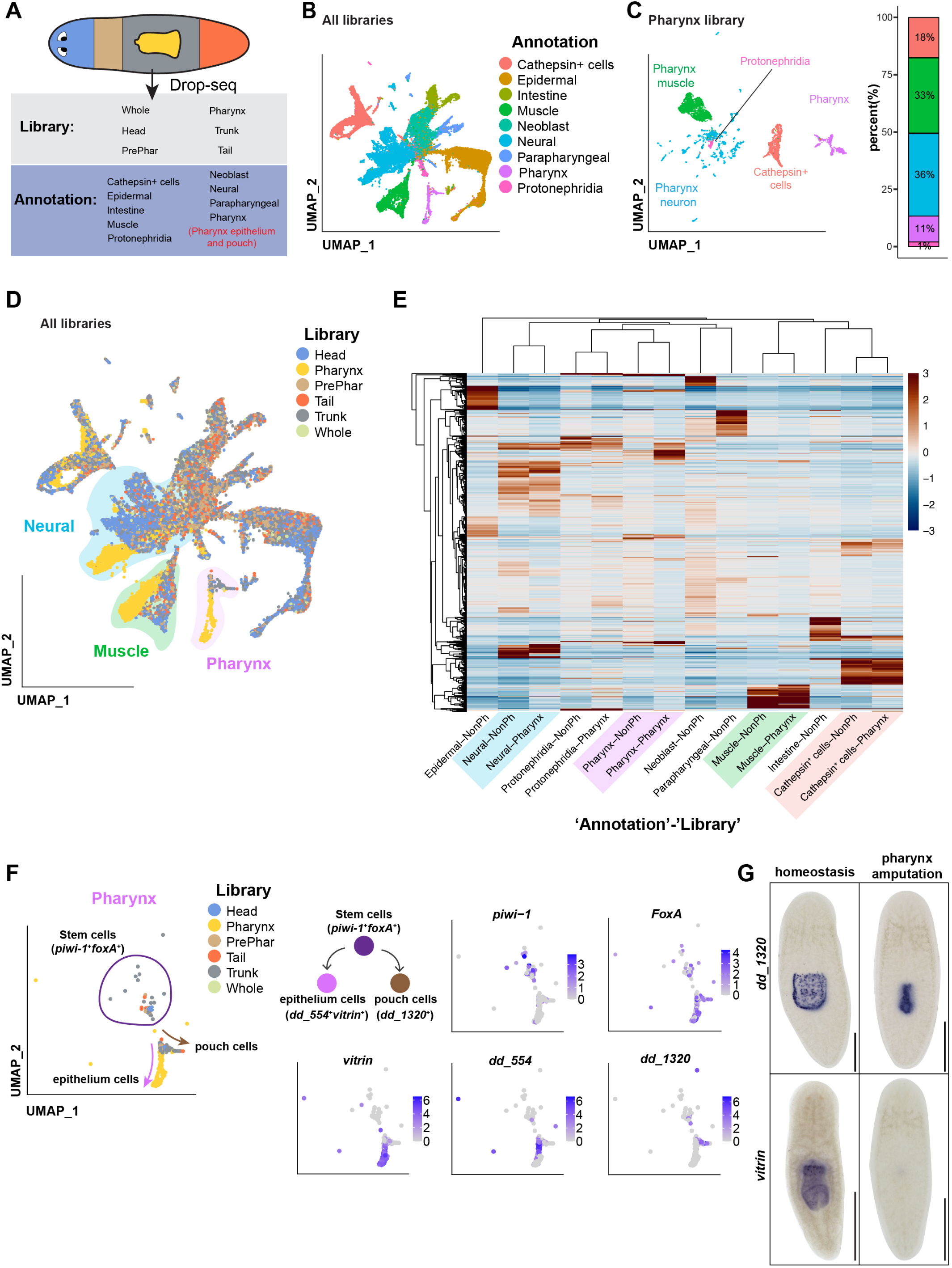
Comprehensive molecular characterization of the planarian pharynx. (A) Schematic overview of region-specific Drop-seq and annotations from Fincher et al., 2018. We propose renaming the original ‘Pharynx’ annotation to ‘Pharynx epithelium and pouch’ (red) to reflect the cell populations more accurately, as discussed in the supplementary text. (B) UMAP plot showing clustering of single-cell transcriptomes from Fincher et al., 2018. Each color represents an annotated cell type. (C) Left: UMAP plot of Pharynx library showing annotated cell types from Fincher et al., 2018. Right: Proportion of major cell types in the pharynx library. (D) UMAP plot of the whole-animal dataset, highlighting specific pharynx cell types (yellow) from Fincher et al., 2018. Subclusters of cells annotated as neural, muscle, and pharynx are shaded. Cells from the pharynx library are adjacent to similar cell types from other body regions. (E) Heatmap of pseudo-bulk transcriptomic profiles comparing cell types as indicated, from either Pharynx or all other libraries (NonPharynx). Columns represent cell populations stratified by both cell-type annotation and the anatomical library of origin (e.g., “Neural-Pharynx,” “Muscle-Pharynx,” etc.), while rows represent differentially expressed genes (1465 genes). Scaled expression: red, higher; blue, lower. (F) Genes in the ‘Pharynx’ cluster represent distinct epithelial cell types. Left: UMAP plot shows the composition of cells from distinct libraries with proposed lineages marked by circle or arrows. Right: UMAP plots of stem cell (*piwi-1*, *foxA*), epithelial (*vitrin*, *dd_554*), and pouch epithelium (*dd_1320*) markers. (G) ISH of *dd_1320* and *vitrin* in homeostatic and pharynx-amputated animals, 1dpa. *vitrin* is absent in pharynx-amputated animals, while *dd_1320* persists, indicating that it marks pouch cells. Scale bars = 500 μm.

**Figure S2.**
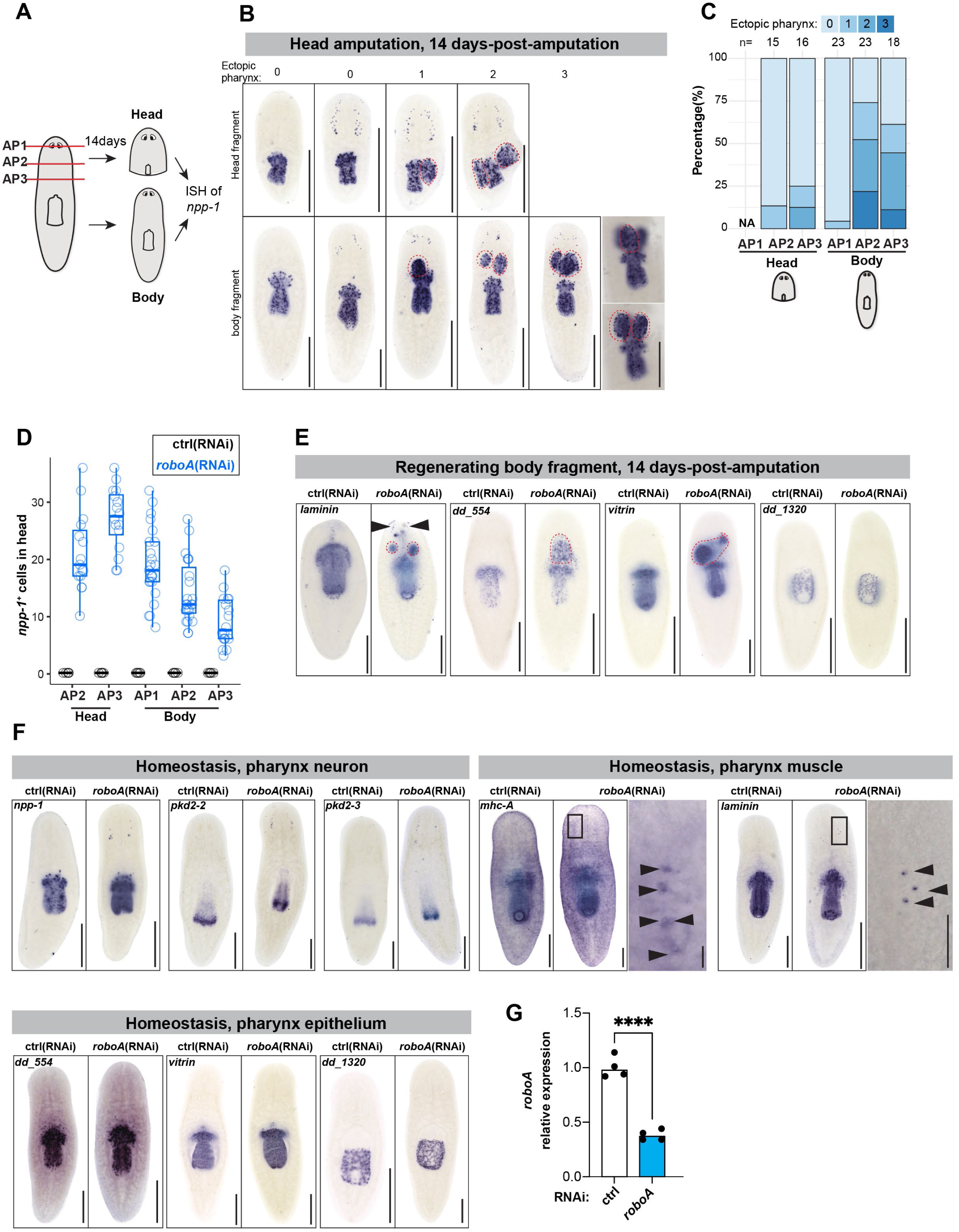
*roboA*(RNAi) induces ectopic pharynges and pharynx cells in the brain. (A) Schematic of experimental design. Head amputation was performed at different anterior-posterior planes (AP1, AP2, AP3). Animals were fixed 14 dpa, followed by ISH for *npp-1*. (B) ISH of *npp-1* in head and body fragments from *roboA*(RNAi) animals 14 dpa. Red dashed circles label ectopic pharynges. Scale bar = 500 μm; inset = 250 μm. (C) Quantification of phenotypes from (B). Removal of more tissue causes higher frequencies of ectopic pharynges. (D) Quantification of EPNs from (B). (E) ISH of pharynx-specific cell types in regenerating body fragments 14 dpa. Ectopic expression of muscle *laminin* (black arrowheads) appears punctate, but other epithelial markers are only observed in ectopic pharynges (red dashed circles). Scale bar = 500 μm. (F) ISH of pharynx-specific cell types in homeostatic animals. Scale bar = 250 μm. (G) qRT-PCR of *roboA* relative to *GAPDH* in RNAi animals, 14dpf. Circles represent individual biological replicates averaged from four technical replicates.

**Figure S3.**
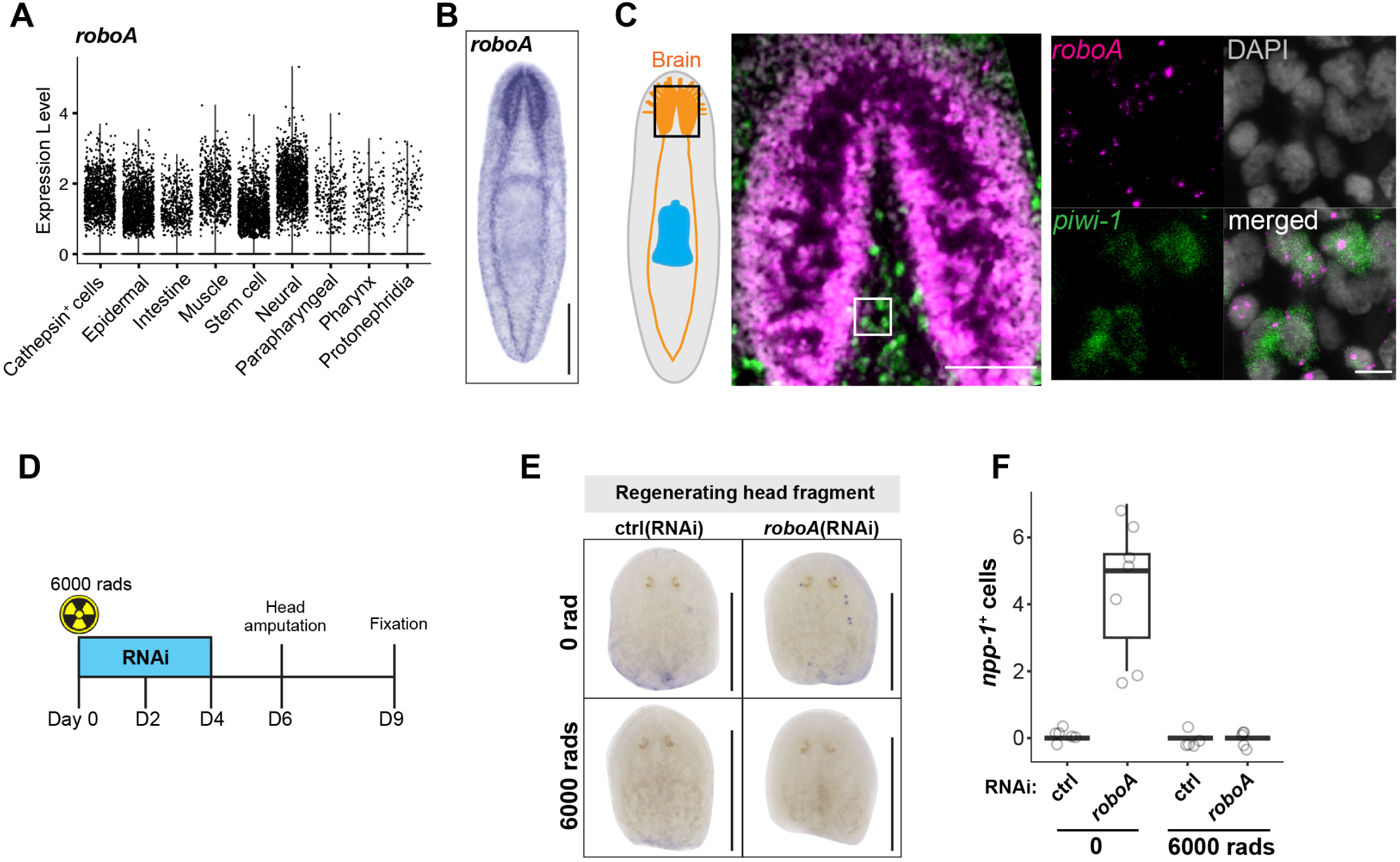
Formation of EPNs requires stem cells. (A) Violin plot showing *roboA* expression in different planarian cell types (derived from Fincher et al., 2018). The y-axis represents normalized UMI counts. (B) ISH of *roboA* in a wildtype animal. Scale bar = 500 μm. (C) Double FISH of *roboA* (magenta) and stem cell marker *piwi-1* (green) in the brain of a homeostatic wildtype animal. Inset is a zoomed image of the white box. Scale bar = 100 µm; inset = 10µm. (D) Schematic of irradiation experiment. Animals were treated with 6000 rads to eliminate stem cells, followed by three feeds of *roboA* RNAi food, decapitated 2 days later, and head fragments were fixed at 3 dpa. (E) ISH of *npp-1* in regenerating head fragments following radiation. Scale bar = 500 μm. (F) Quantification of EPNs in regenerating head fragments from (E).

**Figure S4.**
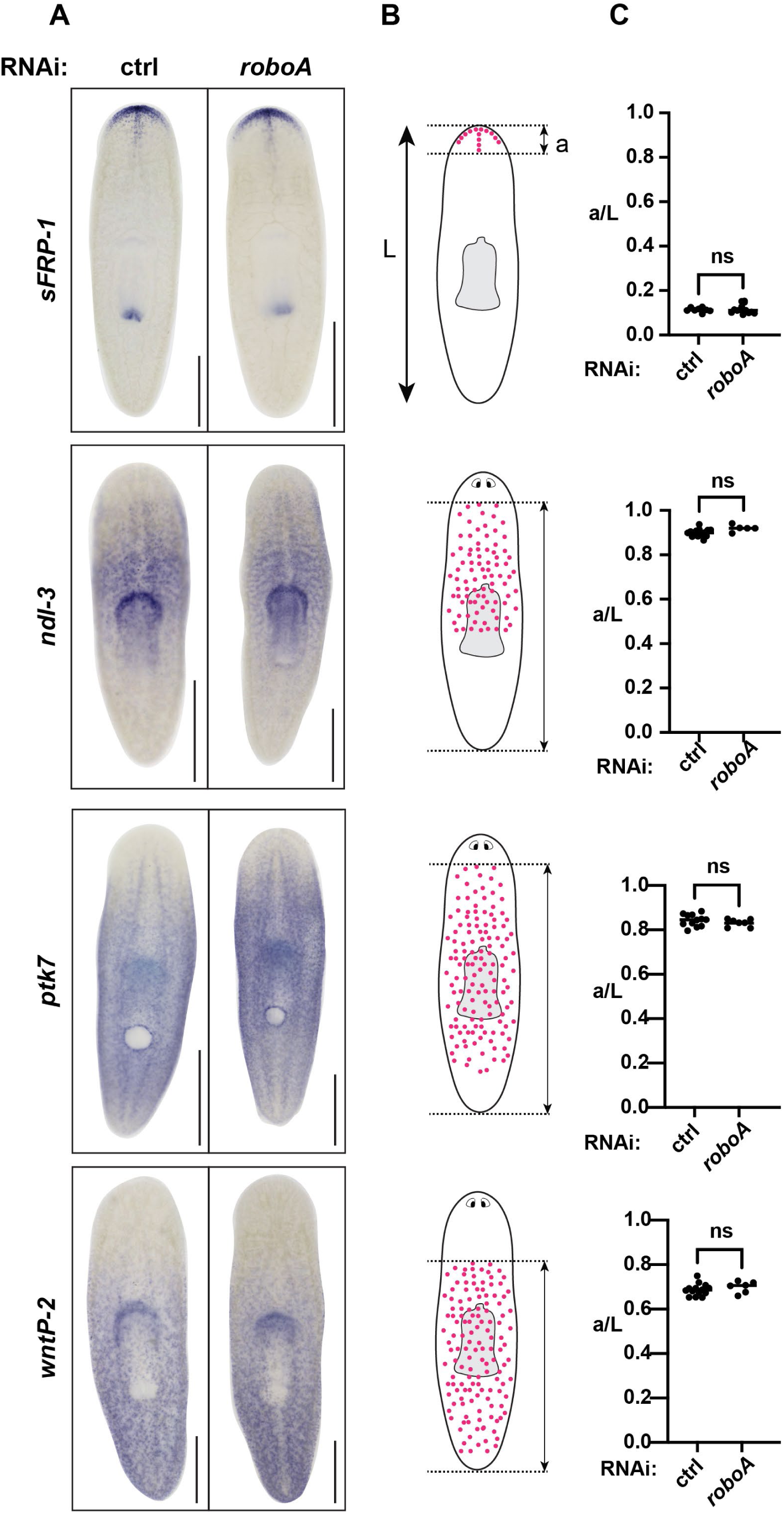
PCG expression is not altered in *roboA*(RNAi) animals. (A) Whole-mount ISH of *sfrp-1, ndl-3, ptk7* and *wnt-P2*. Scale bar = 500 μm. (B) Schematic of measurement strategy. *L* represents the animal length and *a* represents the distance between the anterior and posterior boundaries of the expression domain (between dashed lines). (C) Quantification based on B. Each dot represents an individual animal.

**Figure S5.**
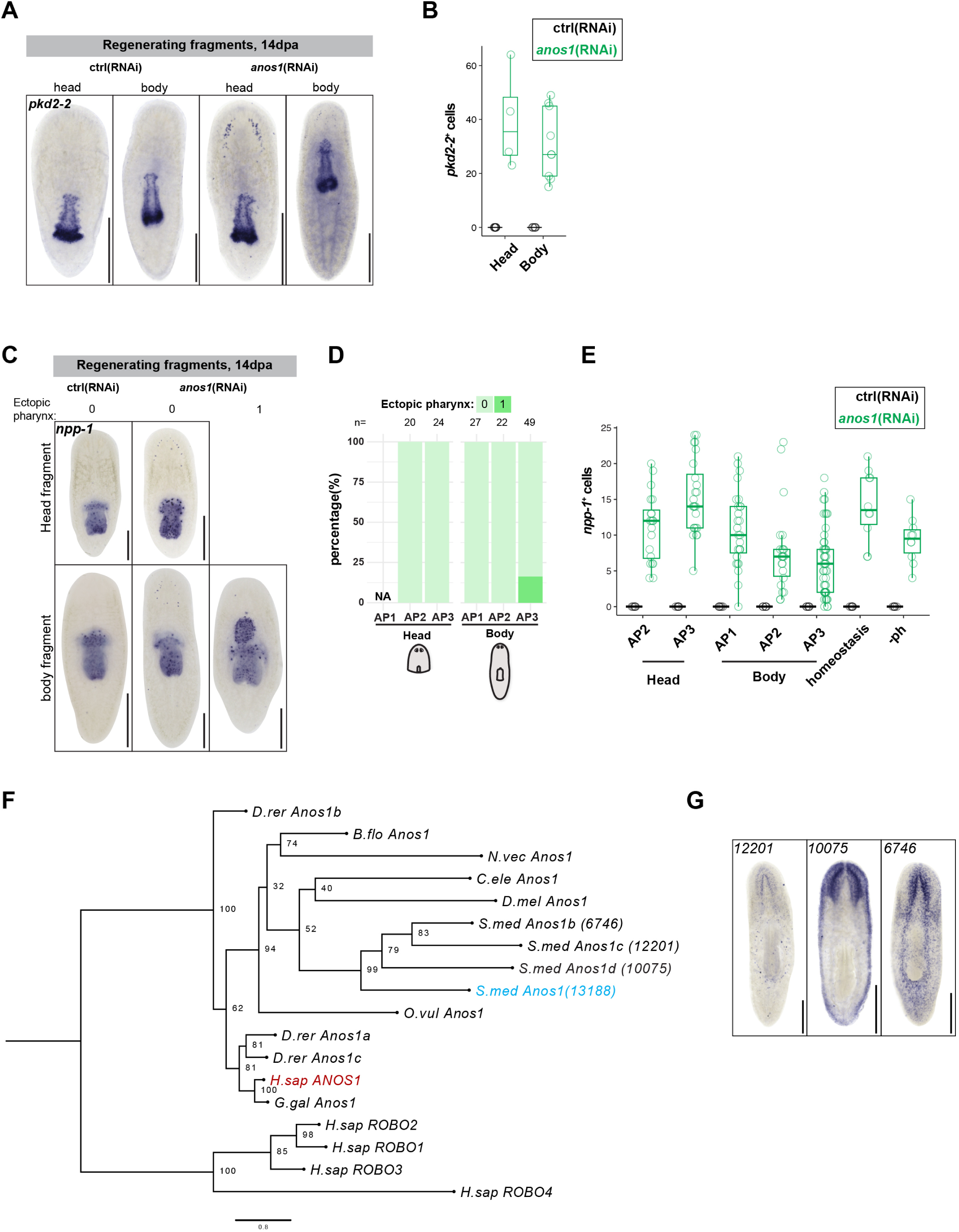
Characterization of *anos1* knockdown phenotype and its phylogeny. (A) ISH of *pkd2-2* in regenerating head and body fragments at 14 dpa. Scale bar = 500 μm. (B) Quantification of ectopic *pkd2-2*^+^ cells in animals from (A). (C) ISH of *npp-1* in regenerating head and body fragments, 14 dpa. Experimental design as in Figure S2A. Scale bar = 500 μm. (D) Quantification of ectopic pharynges in animals from (C). (E) Quantification of EPNs in animals from (C), along with homeostatic and pharynx-amputated animals (-ph). (F) Phylogenetic tree of *anos1* generated by Maximum Likelihood method. The branch length represents the substitution rates. Bootstrap values are labeled on the branch nodes. Accession numbers are in Supplementary Table 3. *S.med, Schmidtea mediterranea; D.rer, Danio rerio; B.flo, Branchiostoma floridae; N.vec, Nematostella vectensis; C.ele, Caenorhabditis elegans; D.mel, Drosophila melanogaster; O.vul, Octopus vulgaris; H.sap, Homo sapiens; G.gal, Gallus gallus*. (G) ISH of *anos1* paralogs in wildtype animals. The numbers represent dd-Smed-v6 transcriptome gene IDs.

**Figure S6.**
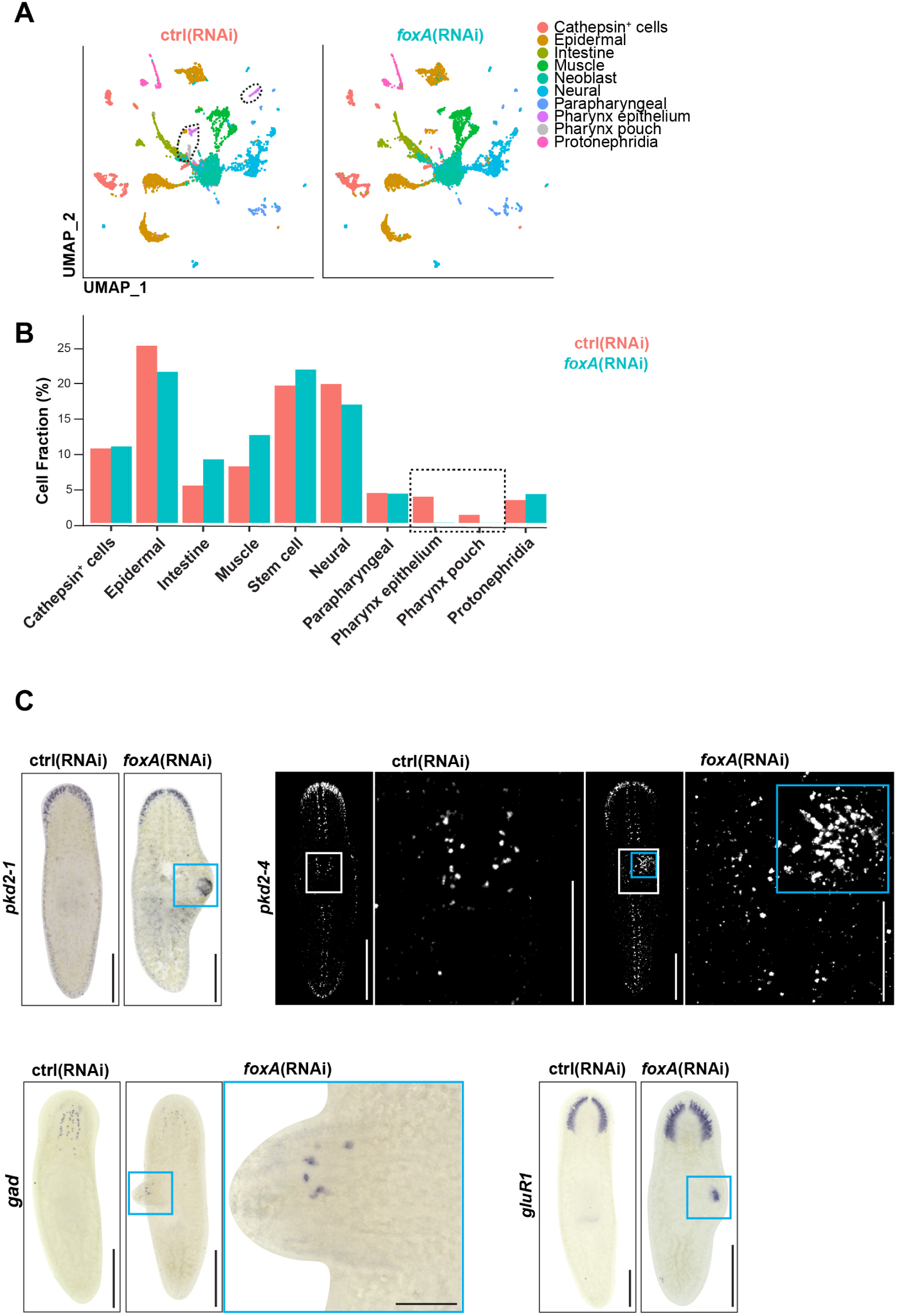
Maintenance of pharynx neuron identity requires *foxA*. (A) UMAP plot of single-cell RNA-seq data from control and *foxA*(RNAi) animals, 21 days after the final administration of RNAi food. Clusters are annotated according to Fincher et al., 2018, along with pharynx-specific annotations as described in Figure S1. Pharynx-specific epithelium and pouch clusters are absent in *foxA*(RNAi) animals. (B) Percentage of cells in annotated clusters in control and *foxA*(RNAi) animals. Dashed black box highlights pharynx epithelium and pouch cells missing in *foxA*(RNAi) animals. (C) ISH of brain-specific neural markers in control and *foxA*(RNAi) animals. Blue box highlights dorsal outgrowth. White box outlines magnified region in inset. Scale bar = 500 μm; inset = 50 μm.

## Notes

https://www.ncbi.nlm.nih.gov/geo/query/acc.cgi?acc=GSE292456

https://github.com/kw572/RoboA

### Competing Interest Statement

The authors have declared no competing interest.

