## Supplementary Text for "Pluripotent Stem Cell Plasticity is Sculpted by a Slit-Independent Robo Pathway in a Regenerative Animal"

**Comprehensive molecular characterization of the planarian pharynx and associated markers**

To systematically define molecular markers of the planarian pharynx, we reanalyzed published single-cell RNA-seq data obtained from dissected anatomical regions ("Libraries") of planarians (Figure S1A) ^1^. Transcriptomic profiles from these libraries were clustered into distinct cell types ("Annotations") based on known marker genes, thus allowing direct correlation between molecular identities and specific anatomical regions (Figure S1B).

Pharynx libraries included cells previously annotated as "Muscle," "Neural," "Cathepsin^+^ cells," "Protonephridia," and "Pharynx" (Figure S1C). This shows a misalignment between annotations and the anatomical libraries because one would expect pharynx libraries to only consist of “Pharynx” cells. Pharynx libraries lacked stem cells ("Neoblasts"), aligning with the fact that the mature pharynx does not contain stem cells. Cells from pharynx libraries annotated as "Muscle," "Neural," and "Cathepsin^+^ cells" clustered closely with cells of the same annotation from other anatomical libraries (Figure S1D). For example, "Neural" cells from the pharynx library clustered closely with "Neural" cells from non-pharynx libraries. Pseudo-bulk transcriptome comparisons further confirmed the similarity of these annotated cell types across libraries (Figure S1E). This analysis shows that the previous annotation accurately captures the unique molecular features of cells from the “Pharynx” library, but preferentially groups neural and muscle cells with non-pharyngeal neural and muscle cells.

In the hierarchical analysis, some cells annotated as “pharynx” were derived from non-pharynx libraries (highlighted in pink) (Figure S1E). To understand what cell types are represented by these ‘pharynx’-annotated cells, we analyzed them using known markers of pharynx cells. The pharynx originates from stem cells expressing *piwi-1* (purple). *foxA* is scattered throughout this cluster, and also expressed in cells negative for *piwi-1*, implying that *foxA*^+^ cells are both stem cells and differentiated cells ^2^ (Figure S1F). Among the differentiated cells, we identified at least two distinct epithelial populations: one branch strongly expressing epithelial markers *vitrin* and *dd_554* (pink), and another branch expressing *dd_1320* (brown) ^1,3^. These two epithelial populations originate from either the pharynx library or a mixture of non-pharynx libraries.

To experimentally validate these putative epithelial markers, we performed *in situ* hybridization (ISH) in homeostatic and pharynx-amputated animals. *Vitrin*, exclusively expressed in pharynx libraries, was completely lost upon pharynx amputation, confirming its specific expression in pharynx epithelial cells ^3^ (Figure S1G). By contrast, *dd_1320* is expressed predominantly in cells from non-pharynx libraries. dd_1320 expression persisted after pharynx amputation, indicating its presence in pharynx pouch cells, a layer of epithelial cells that encircle the pharynx and collapse upon its removal. We conclude that previously annotated “Pharynx” cells include *foxA*^+^ pharynx progenitors, pharynx epithelial cells, and pouch cells. Taken together, we propose revising the previous “Pharynx” annotation to "Pharynx epithelium and pouch" (Figure S1A).
